# RNAJog: Fast Multi-objective RNA Optimization with Autoregressive Reinforcement Learning

**DOI:** 10.1101/2025.08.26.672486

**Authors:** Jiaqi Huang, Ning Feng, Huarong Bai, Yi Fang, Xiaojian Liu, Shengfan Wang, Junchi Yan, Hong-Bin Shen, Zilong Qiu, Ye Yuan, Rongkuan Hu, Xiaoyong Pan

## Abstract

Codon optimization is essential in mRNA vaccine development, while existing tools face limitations in the computational efficiency, sequence diversity and universality. To address these challenges, we develop RNAJog (RNA Joint Optimization with autoregressive Generative model), a framework integrating autoregressive generation with reinforcement learning to optimize codon sequences for minimum free energy (MFE), codon adaptation index (CAI) and GC content, even enabling sequence design without requiring annotated training data. Evaluations in both *in silico* and wet-lab experiments have confirmed RNAJog’s effectiveness and efficiency, with two orders of magnitude faster than traditional algorithm (LinearDesign) for long RNA sequence and about a 10-fold increase in antibody titer compared to the wild-type mRNA for Influenza virus hemagglutinin (HA) mRNA vaccine design in mouse. RNAJog also supports biological constraints for sequence optimization. Using this feature, we minimized m^6^A modification motifs in Bmp2 coding sequence for enhancing the translational efficiency and RNA stability, which are validated in cell-based experiments.

## Introduction

The design and optimization of mRNA sequences is a crucial task in synthetic biology, as it directly impacts the efficiency of protein synthesis and the stability of mRNA in host organisms[1, 2]. Enhanced translation efficiency and structural stability lead to increased protein production in host cells, resulting in improved performance of mRNA-based drug products at lower doses[3]. Moreover, a more stable mRNA structure contributes to greater longevity in vitro, which is essential for the manufacturing and transportation of mRNA therapeutics. A key part of mRNA sequence design is codon optimization. Codon optimization aims to rewrite the coding sequence (CDS) of the mRNA to maximize its expression level and improve the stability, while maintaining the encoded protein sequence. Codon optimization is particularly important in applications such as vaccine development[4], therapeutic protein production[5], etc.

However, codon optimization is a challenging problem due to the degeneracy of the genetic code. Multiple different nucleotide sequences can encode the same amino acid sequence. And selecting the most suitable sequence requires careful consideration of factors such as the translation efficiency and mRNA stability. Existing tools for codon optimization, such as LinearDesign[6], CodonTransformer[7], RiboCode[8], GEMORNA[9], offer valuable approaches, but they have notable limitations. LinearDesign[6] uses a dynamic programming-based strategy to minimize RNA secondary structure free energy, which can be computationally expensive for long sequences. Meanwhile, it can only generate one optimal sequence for a given protein and weight, limiting its diversity and flexibility. CodonTransformer[7], a deep-learning based model, leverages Transformer architecture for context-aware codon optimization. But it focuses solely on codon similarity index (CSI), which may not comprehensively capture the multifaceted objectives involved in codon optimization. Other data-driven methods[8, 10, 11] are mostly constrained by the scale or diversity of their training sets, which poses challenges for practical, real-world applications. For example, RiboCode[8] jointly optimizes MFE and predicted translational efficiency using models trained on Ribo-seq data. While such approaches can perform well within the specific cellular contexts represented in the training data, their generalizability to other endogenous mRNAs across different cell types remains limited[12]. Moreover, some mRNA sequences harbor some biological constraints for better functions in biological processes. For example, N6-Methyladenosine (m^6^A) modification is known to promote the degradation of specific transcripts[13], yet current codon optimization tools have not integrated specific biological subsequence constraints, m^6^A modification motifs and enzyme cleavage sites into their design algorithms. Most available open-sourced codon optimization methods can only manually exclude those subsequences without knowing how to replace them when designing sequences [7, 8].

To address these limitations, we propose the RNAJog model (RNA Joint Optimization with Generative model). RNAJog integrates autoregressive generation with reinforcement learning to optimize codon sequences for multiple objectives simultaneously. Unlike other methods that produce a single sequence or rely heavily on fixed architectures and limited training data, RNAJog efficiently generates diverse, synonymous mRNA sequences. It supports customization of optimization weights and can generate multiple optimized candidates under the same settings, enhancing design flexibility. Furthermore, RNAJog is not constrained by specific training environments and can generalize across different cellular contexts with a known codon frequency table, where RNAJog can be trained on the randomly generated data. We experimentally confirmed RNAJog’s effectiveness with about a 10-fold increase in antibody titer compared to the wild-type mRNA for Influenza virus hemagglutinin (HA) mRNA vaccine design in mouse. Additionally, RNAJog allows for the user-defined biological constraints during sequence optimization, such as enzyme cleavage sites, m^6^A modification constraints, etc. To validate this function for RNAJog, we experimentally validated the designed mRNA sequence of mouse Bmp2 with m^6^A DRACH constraints, demonstrating the improved translational efficiency and RNA stability compared to default optimized and wild-type mRNA sequences.

## Results

### Overview of RNAJog

RNAJog is an autoregressive generative model for optimizing multiple objectives of codon sequences. As shown in Fig.1a, RNAJog consists of three submodules: MFE submodule, CAI submodule, and joint optimization submodule. The MFE submodule generates codon distribution with lower minimum free energy[14], while the CAI submodule generates the distribution with higher codon adaptation index(CAI) [2]. The joint optimization submodule calculates the weighted sum of the codon probability distributions generated by the MFE and CAI submodules and readjust it according to the GC content requirement. Then the model samples from the codon sequence distribution, generating optimized codon sequences that balances all objectives. Fig.1b provides a more detailed illustration of the RNAJog architecture. RNAJog can efficiently generate a large number of synonymous, optimized mRNA sequences. RNAJog allows users to customize the weights of MFE and CAI. And under the same weight settings, RNAJog can generate multiple sequences for selection while LinearDesign only generates one sequence. Given a known codon frequency table, RNAJog can optimize mRNA sequences for any cellular environment, even without requirement to be trained on data specific to that environment. Moreover, as shown in Supplementary Table1, RNAJog has multiple advantages over existing methods, including top-k sequence selection, controllable weight, better extensibility and biological constraint function.

**Fig. 1:**
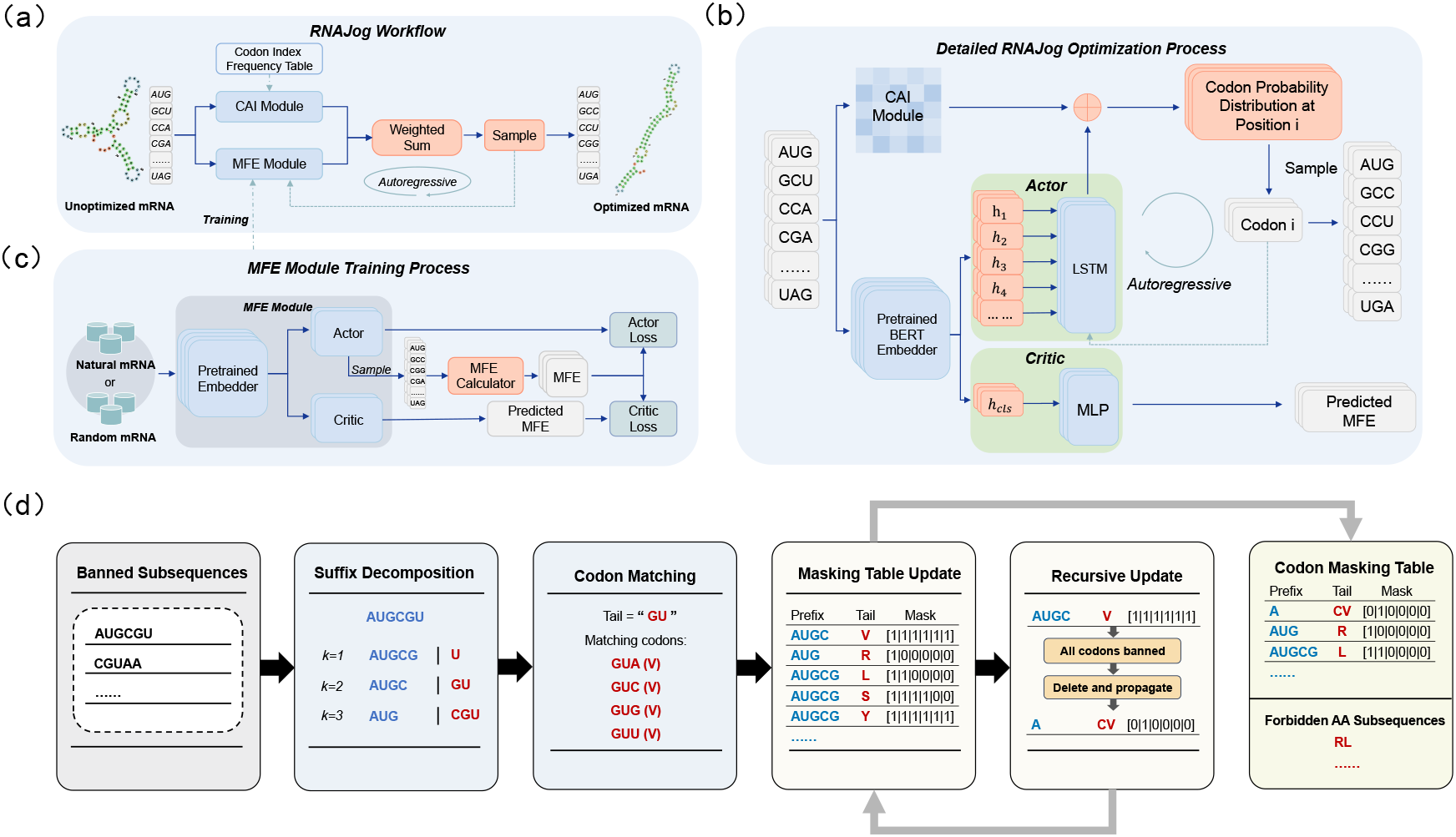
The overview of RNAJog. (a) The RNA sequence generation workflow of RNAJog. (b) The detailed structure of RNAJog model. The model consists of three submodules: MFE, CAI, and joint optimization. The MFE submodule is an Encoder-Decoder model that generates a codon distribution by minimizing the RNA sequence’s MFE. The CAI submodule is fixed and generates a codon distribution by maximizing the CAI. The joint optimization submodule is a sampler that combines the MFE and CAI distributions and outputs a synonymous coding sequence. (c) The training process of MFE submodule. (d) The generation workflow of the codon masking table and the forbidden amino acid subsequences for biological constraints during autoregressive sequence generation.

The MFE submodule is inspired by the Traveling Salesman Problem (TSP) [15]. As shown in Fig.1c, the MFE submodule consists of three main components: a pretrained embedder, an actor, and a critic. The embedder uses the pre-trained language model CodonBERT[16] to encode RNA sequences. The actor, built on Long Short-Term Memory (LSTM), generates codon probability distributions autoregressively based on the embedding and prefix codons. It is trained using policy gradient. The critic is an MLP model that takes the CLS token embedding as input. It is trained using mean squared error, comparing its predictions with the actual MFE of RNA sequences sampled by the actor’s most recent policy. As a value function estimator, the critic helps reduce the variance during training[17].

The MFE submodule is a reinforcement learning model whose supervision signal is derived from the MFE calculated by RNAFold[18]. This method eliminates the need for a training dataset, enabling the model to be trained on randomly generated mRNA sequences. By training from the entire mRNA design space rather that a specific dataset, the model is expected to have better generalization. This could effectively bridge the gap between the limited data availability and robust generalization in the mRNA design problem. The CAI submodule calculates the relative synonymous codon usage (RSUC) of each codon in the RNA sequence and generates the codon probability distribution using the Softmax function. In this way, the submodule can guide the model to generate sequences with higher CAI scores.

In practical applications, there are often design requirements for some biological constraints such as eliminating restriction enzyme sites[19], reducing the number of m^6^A modification sites or other undesirable motifs. While existing design methods generally rely on post-hoc manual mutations to remove these subsequences, such localized patching can disrupt the carefully balanced global properties, often resulting in suboptimal structural stability or codon efficiency. In contrast, integrating these exclusion constraints directly into the optimization phase allows the model to holistically explore the sequence space and identify the best alternative codons, thereby minimizing the compromise in overall performance. To address this challenge, we enhanced RNAJog by incorporating the capability to avoid specific subsequences. As shown in Fig.1d, RNAJog constructs a codon masking table and a forbidden amino acid subsequences list based on the given subsequences to be excluded. Prior to the generation, RNAJog scans the input against the forbidden amino acid subsequences list. If a match is found, it signifies that no mRNA sequence can be constructed that successfully excludes all forbidden nucleotide motifs, as those amino acids inherently necessitate the inclusion of at least one prohibited base-level subsequence. During the autoregressive generation process, the decoder references the codon masking table to mask the prohibited codons and generates the next codon from the remaining valid options.

Moreover, RNAJog is also highly extensible. While MFE and CAI are commonly used metrics in codon optimization, other metrics—such as CSI[20] (Codon Similarity Index)—can be equally important depending on the specific optimization goals. In practice, the landscape of potential metrics varies significantly across different application scenarios that it is practically impossible to pre-define or incorporate every criterion during the initial design of an RNA generation model. In most deep learning–based optimization tools, integrating new metrics typically requires retraining the entire model. However, RNAJog’s probability-based joint optimization framework enables seamless integration of new optimization objectives without the need for complete retraining.

For the experimental design, we trained two different versions of the model, RNAJog and RNAJog-Zero model. RNAJog was trained on the iCodon[21] dataset, which includes four vertebrate transcriptomes: mouse, human, frog, and fish. We used approximately 62,000 RNA sequences from the iCodon dataset for training and reserved 20,000 sequences for evaluation. The other version, RNAJog-Zero, was trained on randomly generated mRNA sequences, and it does not require any annotated training data.

### RNAJog efficiently generates optimized RNA sequences

We first evaluate the computational efficiency of RNAJog compared to LinearDesign. We run RNAJog and LinearDesign for designing RNA sequences with different lengths. As shown in Fig.2a, RNAJog’s codon optimization efficiency is more than two orders of magnitude faster than that of LinearDesign[6], where RNAJog generates sequences using about few seconds even for over 3000-nucleotide sequences. Moreover, RNAJog can generate multiple sequences in a single run for follow-up experimental selection, but LinearDesign generates only one sequence per run. In summary, RNAJog offers a significant advantage in computational speed, especially for long RNA sequences.

**Fig. 2:**
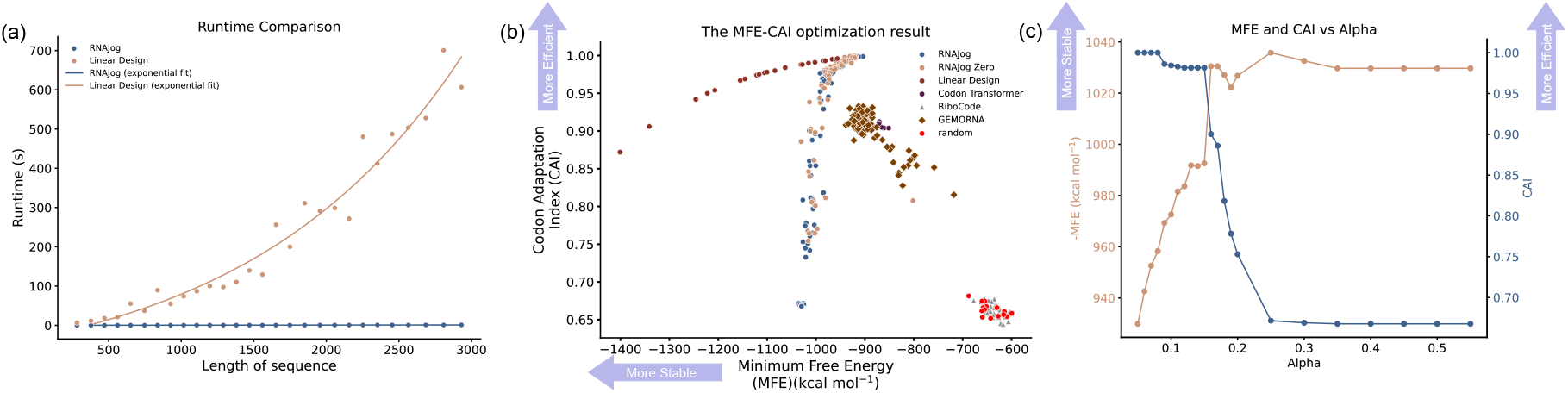
The *in silico* RNA design experiments on natural mRNA dataset. (a)The sequence generation speed for RNAJog and LinearDesign. The runtime comparison between RNAJog and LinearDesign[6]. The computational experiments were conducted on an Intel(R) Xeon(R) Gold 5119T CPU and an NVIDIA GeForce RTX 3090 GPU. The lines in the figure represent the exponential fit of the scatter points. (b) The optimization results for the 2031bp-length mRNA sequence. For all the *in silico* experiments, we use the default settings for these methods, except RiboCode, which is computationally expensive. To reduce the processing time, we run RiboCode with a single iteration per input sequence. random means the sequences are randomly generated. (c) The influence of the MFE-CAI weight alpha on -MFE and CAI (length of CDS: 2031bp). Higher alpha means pay more effort to optimize the CAI of the sequence, lower alpha means pay more effort to optimize the MFE of the sequence. The values on the vertical axis related to MFE in the figure are taken as the negative of MFE, in order to more intuitively illustrate the trade-off between MFE and CAI. The results of other lengths are shown in Supplementary Fig.1.

To assess the optimization effectiveness of RNA sequences across different sequence lengths, we randomly selected six representative RNA sequences (Supplementary Table 2) of varying sizes from the test subset. We evaluated the trained RNAJog model, RNAJog-Zero, LinearDesign[6], CodonTransformer[7], and RiboCode[8] on these sequences. The optimization results on the six representative RNA sequences are visualized using MFE-CAI scatter plots. Fig.2b is the optimization result of the sequence with a length 2031. The optimization results for mRNA sequences of other lengths are shown in the Supplementary Fig.1. As shown in Fig.2b, the RNAJog model outperforms other data-driven codon optimization tools, i.e., CodonTransformer and RiboCode, in optimizing both MFE and CAI simultaneously. RNAJog generates RNAs with similar CAIs and a slightly higher MFE than that of LinearDesign[6], and we can see that RNAJog generates more sequences with high CAIs than that of LinearDesign. In practical applications, a moderate MFE with a high CAI maybe beneficial for RNA translation. On the contrary, LinearDesign tends to generate RNAs with too low MFE, which may hinder the binding or movement of the ribosome, thereby reducing the translation efficiency[3]. Furthermore, we evaluate the influence of the MFE-CAI weight alpha on -MFE and CAI (Fig.2c), the results of other lengths are shown in Supplementary Fig.2. These results show that the MFE-CAI weight parameter, alpha, enables controlled adjustment of the trade-off between MFE and CAI. In addition, RNAJog-Zero, the model trained without any training dataset, also demonstrates the competitive performance compared to RNAJog, demonstrating the strong universality for any cellular environments.

### RNAJog enables *in silico* optimization of SARS-CoV-2 and Cas9 mRNA

We evaluated these models on the SARS-CoV-2 pike protein and compared the results with the commercial mRNA vaccines BNT162b2[22] and mRNA-1273[23]. As shown in Fig.3a, MFE of the commercial mRNA vaccine is close to those of the optimized mRNAs by RNAJog and RNAJog-Zero, and not as low as that of the sequences optimized by LinearDesign[6]. In fact, some studies suggest that too low MFE may reduce protein expression levels due to too stable structure, which is not easy to be open for translation by riobosome[3]. Compared to CodonTransformer and RiboCode, RNAJog offers greater flexibility in adjusting the weighting between MFE and CAI. As shown in Fig.3c and Fig.3d, this controllable weighting allows RNAJog to produce more diverse optimized sequences than CodonTransformer and RiboCode.

**Fig. 3:**
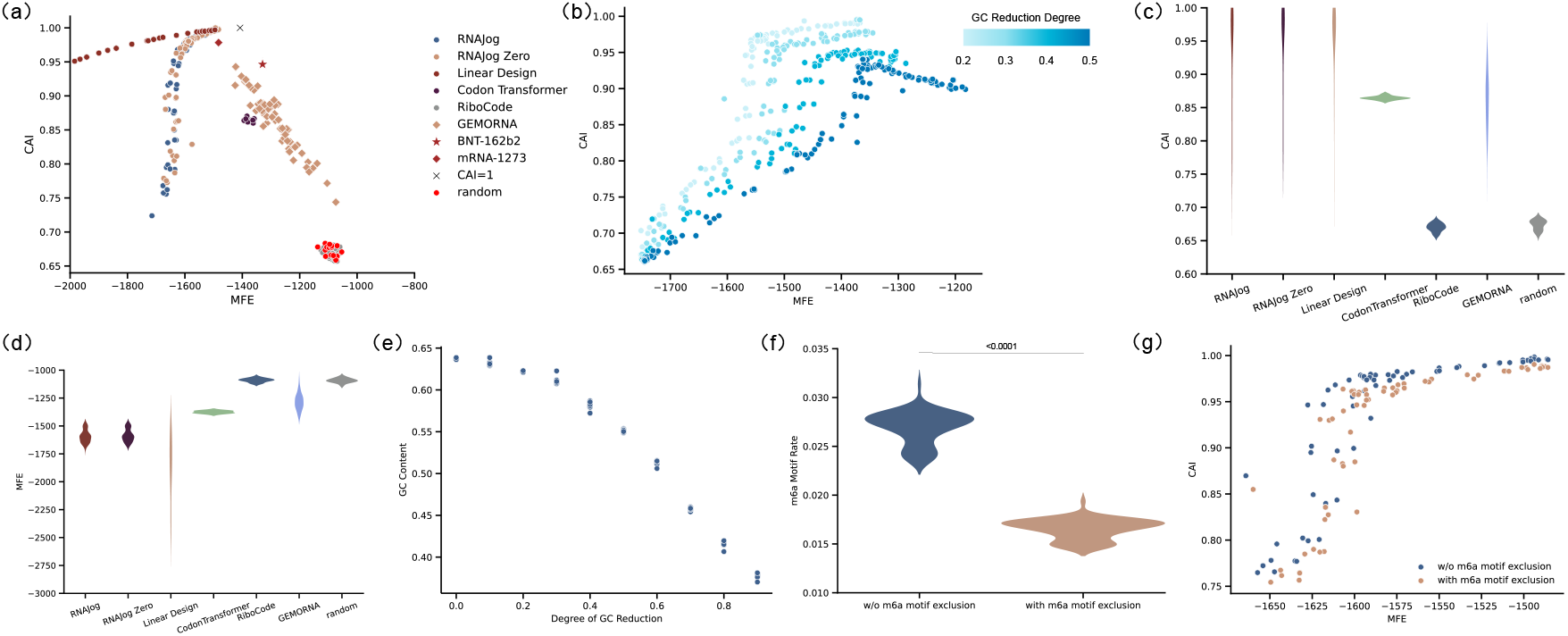
The *in silico* RNA design experiments for SARS-COV-2. (a) The optimization results for the SARS-CoV-2 Spike protein. For all the *in silico* experiments, we use the default settings for these methods, except RiboCode, which is computationally expensive. To reduce the processing time, we run RiboCode with a single iteration per input sequence. random means the sequences are randomly generated. (b) Optimization results under different degrees of GC reduction. A higher GC reduction degree indicates that the model places greater emphasis on reducing GC content rather than optimizing other objectives. (c) The CAI distribution of the SARS-CoV-2 Spike protein optimization results. Adjustable weights lead to a wider CAI distribution. The range of the distribution can be wider for RNAJog, RNAJog-Zero and LinearDesign, which are limited by the test weight range. (d) The MFE distribution of the SARS-CoV-2 Spike protein optimization results. RNAJog and RNAJog-Zero show lower MFE values than CodonTransformer and RiboCode, but still higher than the lower bound of LinearDesign. (e) The influence of GC reduction degree on GC content. (f) Performance of RNAJog in m^6^A motif exclusion. (g) The influence of m^6^A motif exclusion on MFE and CAI of RNAJog.

GC content is another important metric in mRNA design. Imposing a stricter GC reduction constraint effectively decreases the GC content of the optimized mRNAs (Fig.3e), albeit at the cost of a minor trade-off in MFE and CAI (Fig.3b). The relative importance of MFE, CAI and GC content in determining the translation efficiency varies across different contexts. Therefore, generating more diverse and controllable optimized sequences provides mRNA design with a greater flexibility in sequence selection. We also evaluated RNAJog with biological constraint of m^6^A motif exclusion on this protein. As shown in Fig.3f and Fig.3g, RNAJog significantly reduces the occurrence of m^6^A motifs in designed mRNA sequences, with a minimal compromise in MFE and CAI metrics.

To demonstrate the capability of RNAJog for biological constraint of enzyme cleavage sites, we optimize the mRNA sequence of Cas9 with excluding two enzyme cleavage sites XhoI (CTCGAG) and BspQI (GCTCTTC). Under this biological constraint, RNAJog can optimize the mRNA sequence with MFE of −1678.6, and CAI of 0.983, which is improved from the MFE −963.5 and CAI 0.639 of reference mRNA sequence (Supplementary Table 3). However, most methods[7, 8] cannot exclude enzyme cleavage sites during sequence generation. The results demonstrate that the mRNA optimization capability of RNAJog is likely sufficient for most real-world applications.

### RNAJog is experimentally validated for the efficacy of designed HA mRNA vaccines

To showcase the effectiveness of RNAJog, we applied RNAJog and LinearDesign[6] in the real design process of the Influenza virus hemagglutinin (HA) mRNA vaccine with wet-lab experiments. HA is a key protein that mediates virus infecting host cells and the main target of anti-influenza virus neutralizing antibodies, making it the preferred antigen for mRNA vaccine development. Influenza B (B/Washington/02/2019) wild-type (WT) HA Protein was selected for sequence optimization to improve the expression level of HA protein. We designed four mRNA constructs encoding HA proteins (Supplementary Table 4), where HA-M1 and HA-M2 are designed by RNAJog, and HA-M3 and HA-M4 are designed by LinearDesign. The sequence similarity of the four designed mRNAs with the wild-type mRNA are all below 0.7 (Supplementary Table 5). The expression levels of HA proteins in HT1080 and 293T cells transfected with mRNA were detected by flow cytometry (FACS). As shown in Fig.4b and Fig.4c, 24h and 48h after transfection of mRNAs into 293T and HT1080 cells, the tested mRNAs HA-M1, HA-M2 have higher expression levels than that of HA-M3, HA-M4, and the higher expression levels of the four designed mRNAs are higher than that of the mRNA encoding HA-WT protein. 96h and 144h after transfection of mRNAs into 293T cells, the expression levels of HA protein of HA-M1 and HA-M2 were still higher than that of HA-WT, which is higher than that of HA-M3 and HA-M4. Furthermore, the expression levels measured by FACS of five HA proteins in COS7 cells also demonstrate that HA-M1 has better expression levels. Taken together, HA-M1 generally induced the highest expression levels of HA proteins under different experimental conditions.

**Fig. 4:**
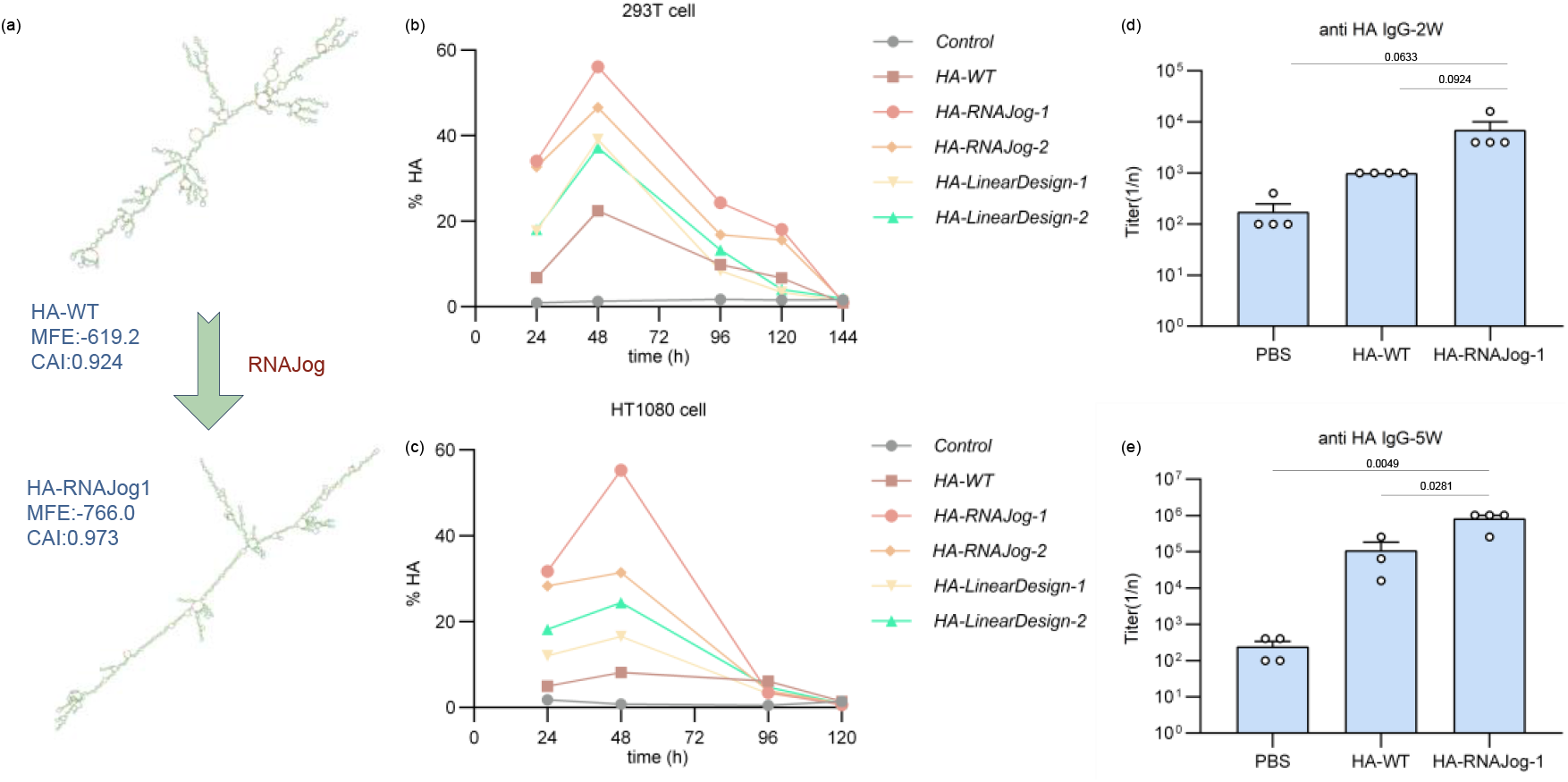
Wet-lab experiments for Influenza virus hemagglutinin (HA) mRNA vaccine design. (a) Secondary structure of the wild-type HA-WT and the optimized HA-M1 by RNAJog. HA-M1 has more hydrogen bonds compared to the unoptimized wild-type, resulting in a lower free energy and a more stable structure. (b) Flow cytometry analyses of 293T transfected with HA mRNAs constructs at different time points, where HA-M1 and HA-M2 are designed by RNAJog, and HA-M3 and HA-M4 are designed by LinearDesign. (c) Flow cytometry analyses of HT1080 cells transfected with HA mRNAs constructs at different time points. (d) HA-binding antibody titers in mouse serum for HA-M1 and HA-WT on day14. (e) HA-binding antibody titers in mouse serum for HA-M1 and HA-WT on day 35. Endpoint titers were defined as the reciprocal of the highest serum dilution with an OD exceeding the cut-off (derived from NC wells). In the HA-WT group, one sample was omitted from the plot because its signal at the starting dilution was below the limit of detection (OD < cut-off).

Furthermore, we investigated the immunogenicity of four mRNA constructs, encapsulated in LNPs STAR0225 in mice. The LNPs (both containing 2 μg mRNA) were used to immunize BALB/c mice (5/female/group) via intramuscular (IM) injections at Day 0 and Day 21. Serum samples were collected on days 14 and 35. The HA-binding antibody IgG titers were analyzed 2 and 5 weeks after the first dose by ELISA. As shown in Fig.4d and Fig.4e, HA-M1-STAR0225 induced high levels of HA-binding antibody IgG titers, with an about 10-fold increase compared to the wild-type HA-WT mRNA. In summary, HA-M1 generally induced the highest expression levels of HA proteins and HA-binding antibody titers, suggesting that RNAJog can be used to improve the expression level in real-world applications.

### RNAJog enables m^6^A modification constraints for differentially regulating Bmp2 expression

N6-Methyladenosine (m^6^A) modification is known to promote the degradation of specific transcripts[13], yet current codon optimization tools have not integrated RNA modification constraints into their design algorithms. To address this gap, we add a biological constraint module into RNAJog, which enables the exclusion of specific subsequences—including m^6^A DRACH motifs—during CDS optimization.

To evaluate the function of RNAjog for m^6^A modification constraints, we selected mouse *Bmp2* (NM_007553), which harboring 30 DRACH motifs within its CDS and exhibited 2.5-fold upregulation upon Mettl3 knockout in the GSE262033 dataset (Supplementary Fig.3a), for wet-lab experiments. We designed Bmp2 mRNA sequences with three RNAJog variants: (1) RNAjog-Bmp2 using default optimization parameters, (2) RNAjog-m6A-1-Bmp2 with automated DRACH motif exclusion, and (3) RNAjog-m6A-2-Bmp2 featuring manually enhanced DRACH removal based on RNAjog-m6A-1-Bmp2. In addition, GEMORNA-optimized Bmp2 with default parameters served as an additional control. All mRNA sequences, along with the predicted MFE and CAI values, are given in Supplementary Table 6.

Following the transfection with equivalent plasmid amounts (Supplementary Fig.3b), RNAjog-Bmp2 markedly elevated Bmp2 protein expression compared to wild-type (Fig.5a and Fig.5b). Intriguingly, while DRACH exclusion in RNAjog-m6A-1-Bmp2 further increased Bmp2 mRNA abundance (RT-qPCR, Fig.5c), the corresponding protein elevation was modest relative to RNAjog-Bmp2 (Fig.5a and Fig.5b), highlighting the complex interplay between m^6^A depletion and translational regulation. Moreover, the expression level of RNAjog-m6A-2-Bmp2, which involved manually enhanced DRACH removal, was lower than that of the RNAjog-m6A-1-Bmp2 (Fig.5b and Fig.5c). This indicates that further manual intervention likely disrupted the global optimization balance achieved by RNAJog, demonstrating that RNAJog already provides a superior, integrated solution that surpasses empirical manual design.

**Fig. 5:**
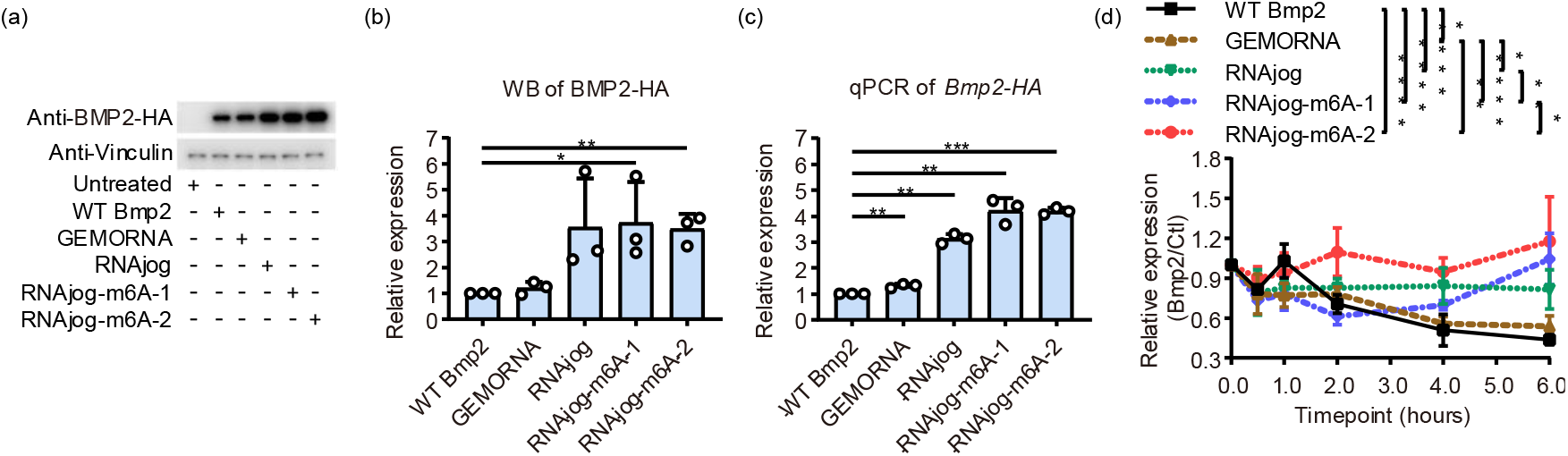
Sequence exclusion of m^6^A motif improve BMP2 mRNA stability and protein production for RNAJog. (a) Representative western blot image showing BMP2-HA detection 48 hours post-transfection with indicated overexpression plasmids or control vectors. Vinculin (VCL) served as a loading control. (b) Western blot quantification. *P*-values were calculated by Student’s t-test. (c) Quantification of *Bmp2-HA* expression after transfection with different plasmids. *P*-values were calculated by Student’s t-test. (d) Wild type and optimized Bmp2 plasmids were transfected into HEK293T cell then treated with 5 mg/mL transcription inhibitor actinomycin D, and collected samples at various time points (0, 0.5, 1, 2, 4 and 6 h) for quantifying the mRNA levels of Bmp2/Ctl. The data are displayed as the means ± SDs. (ANOVA; *p < 0.05, ** p < 0.01, *** p < 0.001, **** p < 0.0001; n = 3).

To investigate mRNA stability during codon optimization. We analyzed mRNA stability in RNAjog (standard codon-optimized) and RNAjog-m6A-1/2 (m^6^A motif sequence-exclusion) compared with their wild-type and GEMORNA counterparts. Notably, RNAjog exhibited significantly increased mRNA relative to both wild-type and GEMORNA controls. Furthermore, after using RNAJog with m^6^A avoidance model, mRNA stability was further increased (Fig.5d).

## Discussion

In this study, we developed RNAJog, a framework that integrates autoregressive sequence generation with reinforcement learning to address the high-dimensional search space with biological constraints and multi-objective optimization challenges in mRNA design. The *in silico* and wet-lab experimental results demonstrate that RNAJog outperforms existing data-driven approaches in terms of the optimization quality, computational efficiency, and sequence diversity.

Despite these advantages, several limitations remain. First, the optimization objectives in RNAJog rely on computational metrics such as minimum free energy (MFE), codon adaptation index (CAI), and GC content, which serve as proxies for mRNA stability and translation efficiency. While these metrics enable generalization across diverse contexts, they are not strictly predictive of biological outcomes, and objective weights may need to be empirically tuned based on experimental feedback. Second, important biological factors such as co-translational protein folding are not modeled. Previous studies[24] suggest that excessively rapid translation may impair proper protein folding; however, due to the lack of large-scale, high-quality datasets, protein misfolding risk is not currently incorporated into RNAJog framework. Third, RNAJog focuses on the coding sequence (CDS) optimization and does not consider untranslated regions (UTRs), which play critical roles in post-transcriptional regulation. Joint optimization of CDS and UTR sequences represents a critical next step toward improving biological fidelity.

Finally, we have open-sourced RNAJog, including modules for excluding specific subsequences, to facilitate adoption by the research community. Its modular and extensible architecture enables seamless integration of additional optimization objectives. Future work will focus on incorporating new optimization modules and integrating RNAJog with UTR design models to establish a comprehensive end-to-end mRNA design framework suitable for real-world applications.

## Methods

In this study, we introduce RNAJog, an RNA optimization method that combines autoregressive generation with reinforcement learning to simultaneously optimize both the minimum free energy (MFE) and codon adaptation index (CAI) Furthermore, RNAJog can be trained on randomly generated dataset, enhancing its adaptability and practical applicability across diverse cellular environments. Finally, we experimentally verified the effectiveness of RNAJog for HA mRNA vaccine design and Bmp2 with m6A modification constraints.

### Problem statement of RNA design

In general, the problem of codon optimization can be described as a combinational optimization problem[6]. For a given protein sequence *p* = (*p*_1_, *p*_2_, *p*_3_, …, *p*_*n*_), we try to find a codon sequence *c* = (*c*_1_, *c*_2_, *c*_3_, …, *c*_*n*_) that minimize the cost function *F*(*c*), where *c* is the codon sequence of *p*. The cost function *F* is defined as a weighted sum of the negative of codon adaptation index and the minimum free energy of the RNA sequence.

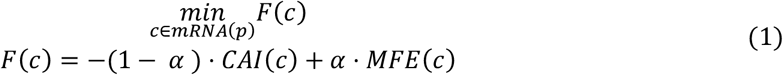

where *α* is pre-defined weight parameter.

### Model architecture of RNAJog

The RNAJog model is a generative model which generates the optimized CDS (coding sequence) of RNA with a given protein or RNA CDS, as shown in Fig.1a and a more detailed structure of RNAJog is given in Fig.1c. The model can be divided into 3 parts: MFE submodule, CAI submodule and joint optimization sFubmodule. The MFE submodule is an Encoder-Decoder model that generates the codon distribution of the RNA sequence with a lower MFE. The CAI submodule is an untrainable module that generates the codon distribution of the RNA sequence with a higher CAI. The joint optimization submodule is a sampler, which samples from the weighted sum of the codon distribution generated by the MFE submodule and CAI submodule. The output of the joint optimization submodule is a synonymous coding sequence of the input sequence. RNAJog jointly optimizes the MFE and CAI values, and it can be easily extended to cover more properties of RNAs.

#### CAI submodule

According to the definition, the CAI of a codon sequence is the geometric mean of the relative synonymous codon usage (RSUC) of each codon in the sequence *c* = (*c*_1_, *c*_2_, *c*_3_, …, *c*_*n*_).

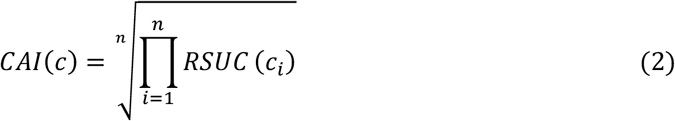

The RSUC of a codon is defined as the frequency of the codon in the codon pool of the corresponding amino acid.

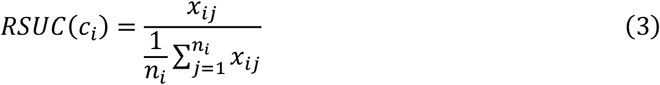

where *x*_*ij*_ is the frequency of the codon *c*_*i*_ in the codon pool of the amino acid *a*_*j*_, *n*_*i*_ is the number of codons that encode the amino acid *a*_*i*_.

Since the frequency of codons varies in different species, the RSUC of a codon is different across different species. Before calculating the RSUC of a codon, we need to input the codon usage table of the species to the CAI submodule. The RSUC of each codon could be seen as a score. We can use the score to generate the codon distribution in which the codons with a higher RSUC have a higher probability to be sampled. Specifically, we can use the Softmax function to calculate the probability of each codon.

The CAI submodule takes the RNA sequence as input. First, it calculates the RSUC of each codon in the sequence. Then, it calculates the probability of each codon using the Softmax function. The output of the CAI submodule is the codon probability distribution of the RNA sequence.

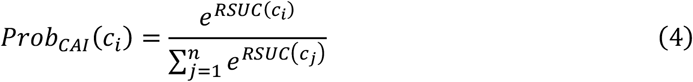

### MFE Submodule based on an Actor-Critic model

The MFE submodule is an Actor-Critic model that generates the codon distribution of the RNA sequence with a lower MFE. The MFE submodule takes the RNA sequence as input, as shown in Fig.1b. Firstly, we use the pre-trained CodonBERT[16] model to embed the RNA sequence into an embedding tensor of a shape (n+1, 768), where n is the num of codons. The additional vector in the embedding tensor corresponding to the CLS token, which is used as the representation of the whole sequence. Then, we use the actor to generate the optimized RNA sequence.

### Autoregressive actor based on an LSTM

The actor is an LSTM model that generates the codon distribution of the RNA sequence with the embedding tensor of the RNA sequence as the input. The output of the actor is the probability distribution of each codon, where the sequence is generated in an autoregressive manner. The probability distribution of the *i*-th codon is calculated based on the prefix codons and the *i*-th token embedding in the embedding tensor.

### Critic with a multilayer perceptron

The critic in this model is a multilayer perceptron (MLP) that processes the embedding of the CLS token as input. Its primary role is to estimate the expected MFE of RNA sequences sampled by the actor’s most recent policy. To achieve this, the critic is trained using a mean squared error (MSE) loss, where it learns to minimize the difference between its predicted MFE values and the actual MFE of RNA sequences sampled under the most recent policy of the actor. By providing baseline to the actor, the critic helps stabilize training and reduce variance, leading to more efficient learning.

### Training Process of the MFE Submodule

We use the policy gradient to train the MFE submodule[15], as the pseudocode shown in Algorithm 1.*S* represents the training set containing sequences, *T* denotes the number of training steps, and *B* is the batch size. *θ*_*actor*_ and *θ*_*critic*_ are the parameters of the Actor and Critic networks, respectively. *s*^*i*^ is a sequence sampled from *S, c*^*i*^ represents codons sampled from the Actor network given *s*^*i*^, and *b*_*i*_ is the baseline value estimated by the Critic network for *s*^*i*^. 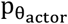is the stochastic policy of actor with parameters *θ*_*actor*_. *MFE*_*i*_ is the Minimum Free Energy computed by RNAFold for *c*^*i*^. *Loss*_*actor*_ is the Actor loss, while *Loss*_*critic*_ is the Critic loss. *α* is the weighting factor for the Critic loss. ADAM is the optimization algorithm used to update *θ*_*actor*_ and *θ*_*critic*_.

#### Algorithm 1

Training process of MFE Submodule

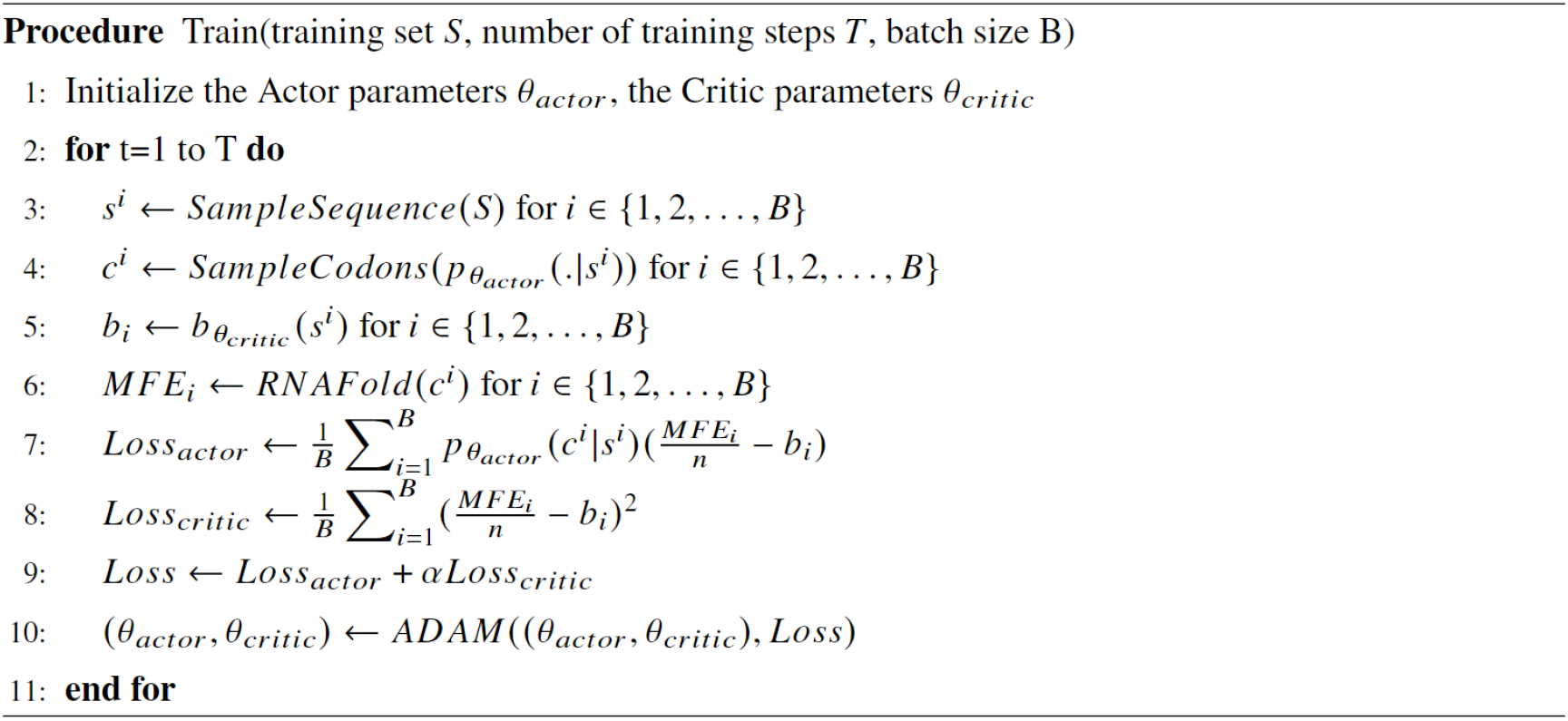

The final loss function of the MFE submodule is defined as follows:

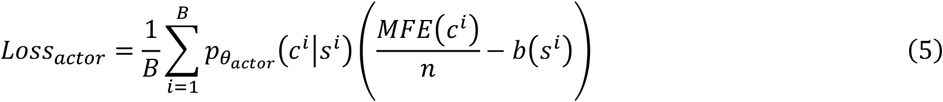

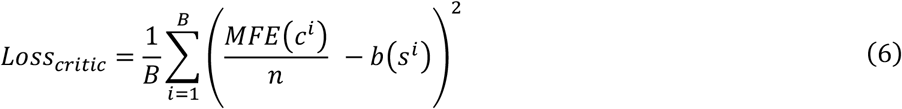

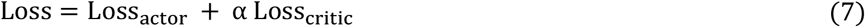

where B is the number of sequences,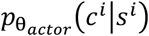is the conditional probability of the codon sequence *c*^*i*^ with given input *s*^*i*^,n is the number of codons in *c*^*i*^. *MFE*(*c*^*i*^) denotes the actual MFE of the RNA sequence *c*^*i*^. *b*(*s*^*i*^) denotes the output of critic. The *Loss* is composed of two components: the actor loss *Loss*_*actor*_ and the critic loss *Loss*_*critic*_. α is the hyperparameter that balances the actor loss and the critic loss.

To train the RNAJog, we first calculate the actual MFE of the optimized codon sequence using RNAfold[18]. Next, we define the advantage of c as 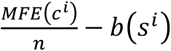. This advantage measures how good the current sequence is compared to the expected performance of the actor. Advantage serves as the reward in the policy gradient training process. Finally we use the advantage and the log probability to update the parameters of the MFE actor submodule with the policy gradient method[17].

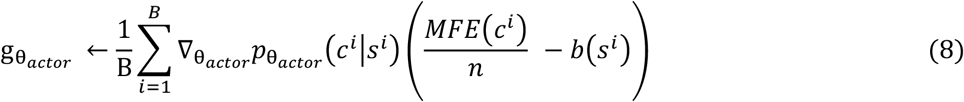

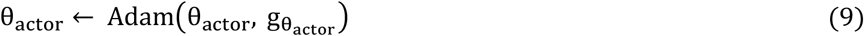

### Generation of the random data for the training of RNAJog-Zero

The training data of RNAJog-Zero is randomly generated using the following approach. First, protein sequences of varying lengths are randomly generated, with each amino acid sampled uniformly from the set of 20 standard amino acids. Next, each amino acid in the protein sequence is translated into its corresponding codon. For amino acids that are encoded by multiple synonymous codons, one codon is randomly selected. Finally, the start codon AUG is appended to the beginning of the codon sequence, and a stop codon, randomly chosen from UAA, UAG, or UGA, is appended to the end of the sequence.

### Designing mRNA sequence of mouse Bmp2 using RNAJog with m6A DRACH constraints

To evaluate the practical efficacy of biological m6A constraints, we applied RNAJog to design mouse Bmp2 mRNA while specifically depleting m^6^A-associated DRACH motifs. We maintained a prioritized list of DRACH motifs, ranked by their reported association with m^6^A modification. This comprehensive list was initially provided to the model as a set of banned subsequences. Recognizing that certain amino acid sequences may inherently require specific motifs, we implemented an iterative optimization strategy: if the model determines that a valid mRNA sequence cannot be generated under the current stringent constraints, it automatically removes the motif with the lowest m^6^A-association from the forbidden amino acid subsequences list and re-attempts the generation process. This step-wise refinement continues until a successful candidate is identified, ensuring that the final sequence achieves the maximum possible reduction of m^6^A motifs while maintaining strict adherence to the target protein sequence.

### Wet-lab experiments

#### Expression of mRNA in vitro

HT1080 and 293T cells were inoculated in 24-well plates and cultured in DMEM medium (HyClone, SH30022.01) supplemented with 10% FBS (HyClone, SH30406.05) at 37°C in 5% CO_2_ for 24 hours. The cells were transfected with mRNA of HA protein using Lipofectamine 2000 (Invitrogen). Cells were collected after transfection. The expression of HA protein was then detected by flow cytometry with Influenza B Hemagglutinin/HA Antibody, Rabbit MAb (Sino Biological, 11053-R004) and a secondary antibody Goat Anti-Rabbit IgG H&L (Alexa Fluor^®^ 488) (Abcam, ab6717).

#### Preparation of mRNA

In vitro transcription was performed using the TranscriptAid T7 High Yield Transcription Kit (Thermo K0441), using linear DNA as a template for generating mRNA, with addition of a certain proportion of pseudouridine and capping reagents. In vitro transcription conditions: the reaction system was configured according to the kit instructions, reaction at 37 °C for 0.5-2 hours, the transcript was digested for 30 minutes by DNase, and the transcript was purified with Monarch RNA Cleanup Kit (NEB T2040L).

### Lipid nanoparticle preparation

mRNA was dissolved in citrate buffer (pH 4) and adjusted the concentration of mRNA to 0.2 mg/mL, so that obtaining aqueous layer thereby. A1-EP10-O18A (synthesized according to our patented method), 1,2-distearoyl-sn-glycero-3-phosphocholine (DSPC), cholesterol and DMG-PEG2000 were dissolved in desired molar fractions in dry ethanol and the total concentration of lipids was adjusted to 10 mg/mL, thereby obtaining organic layer. The aqueous layer and organic layer were admixed in 3:1 ratio (v/v) by microfluidic device (NanoAssemblr® Ignite™) at total flow rate of 12 mL/min. The mixture was 10-fold diluted with PBS buffer (pH 7.4). Ethanol was separated by tangential flow filtration (Repligen, TFF). The solution was concentrated to 0.1 mg/mL (mRNA concentration) and filtrated by 0.22 μm millipore filter to afford mRNA containing lipid nanoparticles.

#### Mouse immunizations

6-8 weeks old female BALB/c mice(4/group) were immunized with PBS or test vaccine by intramuscular injections on day 0 and day 21. Serum samples were collected on days 14 and day 35 for detection of anti HA IgG antibody titers.

#### ELISA

ELISA plates were coated with 1μg /mL Influenza B (B/Washington/02/2019) Hemagglutinin / HA Protein (His Tag) (Sino Biological, 40722-V08H) overnight at 4°C. The coated plates were washed and incubated with a given serum dilution. Anti-mouse antibodies labeled with HRP (Abcam, ab6728.) were used to measure the binding of antibodies specifically to HA protein using TMB substrate. The absorbance values at 450 nm were measured using a multi-functional microplate reader, and the anti HA IgG antibody titers of serum samples of each animal were analyzed using a four-parameter fitting method.

#### Fluorescence-Activated Cell Sorting (FACS)

24 h,48h,96h,120h and 144h after transfection, the supernatants of the cells were discarded. And cells were digested with 0.25% trypsin for 1 minute and centrifuged at 1500 rpm for 5 minutes. The supernatants were discarded. Then the cell pellets were suspended with FACS buffer (PBS containing 2%FBS) and centrifuged at 1500 rpm for 3 minutes. Repeat washing step twice. Influenza B Hemagglutinin/HA Antibody were used for staining. Add the above antibodies to cells and incubate at 4°C for 1 h. After repeat washing step twice, add Goat Anti-Rabbit IgG H&L (Alexa Fluor® 488) antibodies to cells and incubate at 4°C for 1 h. The cells were washed twice and finally re-suspended with 300μL with FACS buffer. The analysis was performed by flow cytometry (FACS). Flow cytometric data were quantitatively evaluated using FlowJo software.

#### Plasmid Generation

The coding sequence (CDS) of mouse Bmp2 (NM_007553) was codon-optimized using GEMORNA[9] or RNAjog software. The wild type and optimized CDS was fused in-frame with a glycine-serine (GS) linker and a C-terminal hemagglutinin (HA) tag. The resulting fusion construct was subcloned into the pcDNA3.1(+) expression vector between KpnI and BamHI restriction sites within the multiple cloning site. Plasmid assembly was performed as a commercial service by Sangon Biotech (Shanghai, China).

#### Cell Culture and transfection

HEK293T was cultured in a humidified incubator at 37°C (5% CO2) in Dulbecco’s Modified Eagle Medium (DMEM-Glutamax) that was supplemented with 10% Fetal Bovine Serum (FBS) (both Thermo Fisher Scientific) and 1% Penicillin, Streptomycin and Amphotericin B (PSA). For Lipofectamine 2000 transfection, transfection was as according to manufacturer’s instructions. A ratio of 1 µl of Lipofectamine 2000 to 1 µg of plasmid DNA was used. A total of 500 ng plasmid DNA was used for 24-well plates.

#### WB

After 48 hours transfection, cells were washed with PBS, and then lysed in RIPA buffer (Beyotime Biotechnology) with Protease inhibitor cocktail (Beyotime Biotechnology). The lysates were cleared by centrifugation at 12,000 g for 10 minutes at 4°C. Protein concentrations were determined by BCA Protein Assay (Beyotime Biotechnology) according to manufacturer’s instructions. Equal amounts of total protein were loaded onto precast 4-20% Tris-Gly mini gels (Shanghai Wansheng Haotian Biotechnology) and separated by sodium dodecyl sulfate – polyacrylamide gel electrophoresis (SDS-PAGE). Protein samples were electrotransfered onto a 0.45 µm PVDF (polyvinylidene fluoride) membrane (Merck Millipore). Membranes were blocked by 5% bovine serum albumin (BSA) supplemented with Tween 20 (TBST). Blocked membranes were incubated with primary antibodies overnight in 2% BSA in TBST at 4°C. The next day, membranes were washed three times with TBST (10 minutes each), followed by incubation with secondary antibodies for 1 hour at room temperature. Membranes were washed three times and signal developed using Pierce ECL Western Substrate (Thermo Fisher Scientific). Blots were visualized using the ChemiDoc MP Imaging System. Details of antibodies are provided in Supplementary Table 7.

#### RT-qPCR

To quantify overexpressed *Bmp2* transcripts, total RNA was extracted by the TRIzol reagent (Thermo Fisher Scientific). Reverse transcription was performed using the PrimeScript RT reagent Kit with gDNA Eraser (Perfect Real Time) (Takara) as according to manufacturer’s instructions using 200-1000 ng (typically 400 ng) input total RNA. cDNA was diluted 1:5 in nuclease-free water prior to quantitative polymerase chain reaction (qPCR) amplification using 2X Universal SYBR Green Fast qPCR Mix (ABclonal Biotechnology) and the QuantStudio 5 Real-Time PCR System. Universal cycling conditions were used (i.e. 95°C for 10 minutes, followed by 40 cycles of 95°C for 15 seconds and 65°C for 1 minute). 10 pmols of each primer were included per reaction. Reaction specificity was confirmed by post-run melting curve analysis. Primer sequences are listed in Supplementary Table 8.

#### mRNA stability assay

Cells were transfected with the indicated plasmids and treated with 5 μg/ml Actinomycin D (HY-17559, MCE) to block transcription. Total RNA was harvested at the indicated time points (0, 0.5, 1, 2, 4, and 6 hours) using Trizol reagent (Thermo Fisher Scientific) and reverse-transcribed into cDNA with a PrimeScript RT kit (RR047A, Takara). cDNA was subjected to RT-qPCR analysis as previously described. Primer sequences are listed in Supplementary Table 8. An SV40 promoter-driven gene encoded in the plasmid was used as an internal control (Ctl). Data represent three independent experiments.

## Statistical Analysis

For comparisons of two samples, a Student’s t-test was used. For comparisons of more than two groups, an ordinary one-way analysis of variance (ANOVA) was performed with Bonferroni’s post hoc test for inter-group comparisons.

## Data availability

We used iCodon dataset (www.iCodon.org) for training RNAJog model. The processed dataset and experimental data are available at http://www.csbio.sjtu.edu.cn/bioinf2/RNAJog/Data.htm

## Code availability

Source codes are available at https://github.com/kxstd/RNAJog. To facilitate the use of RNAJog, we developed a user-friendly web server at http://www.csbio.sjtu.edu.cn/bioinf2/RNAJog/.

## Contributions

X.P., R.H., Y. Y. and Z.Q. conceptualized the project. J.H. was responsible for dataset collection and preprocessing. The model was developed by J.H., with methodological guidance provided by J.Y. and X.P., and J.H. carried out model training. X.P. and H.B.S. supervised and provided strategic direction for the project. Y.F. developed the online website and implemented its functionality. N.F. performed the sequence selection and subsequent optimization of Bmp2 mRNA variants and experimentally validated the efficacy of RNAJog in designing m^6^A-motif-constrained sequences. H.B. and R.H. experimentally verified the effectiveness of RNAJog for HA vaccine design, with contributions from Y.Y., J.H., H.B., X.P. drafted the manuscript, with contributions from X.L. and S.W., and received constructive feedback from H.B.S. All authors reviewed and approved the final manuscript.

