## Supplementary Information for "RNAJog: Fast Multi-objective RNA Optimization with Autoregressive Reinforcement Learning"

### Supplementary Material

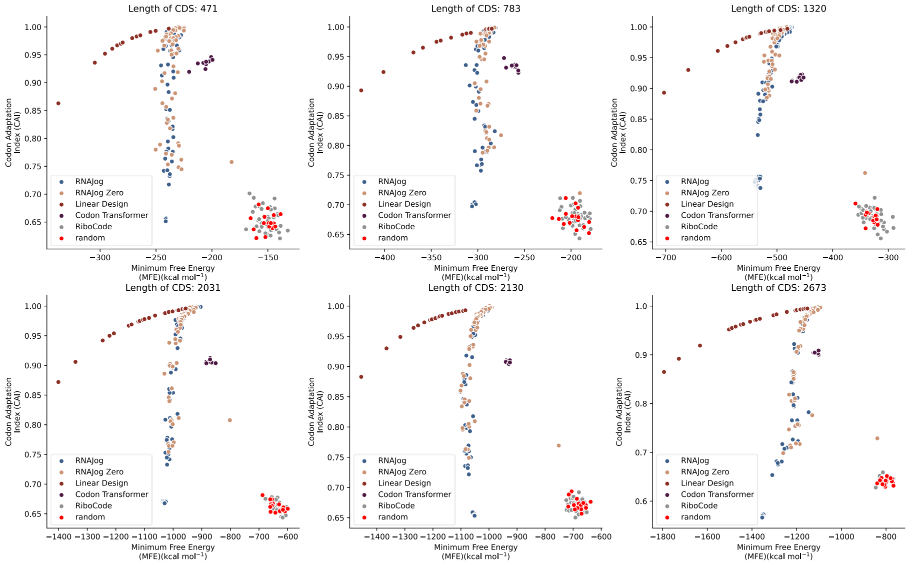


**Supplementary Figure 1.** The MFE–CAI optimization results of different methods on six mRNA sequences with different lengths from 471 to 2673. RNAJog, RNAJog-Zero and LinearDesign can set the weight parameters between MFE and CAI. For all the in silico experiments, we use the default settings for these methods, except for RiboCode, which is computationally expensive. To reduce the processing time, we run RiboCode with a single iteration per input sequence. Here random means the sequences are randomly generated.

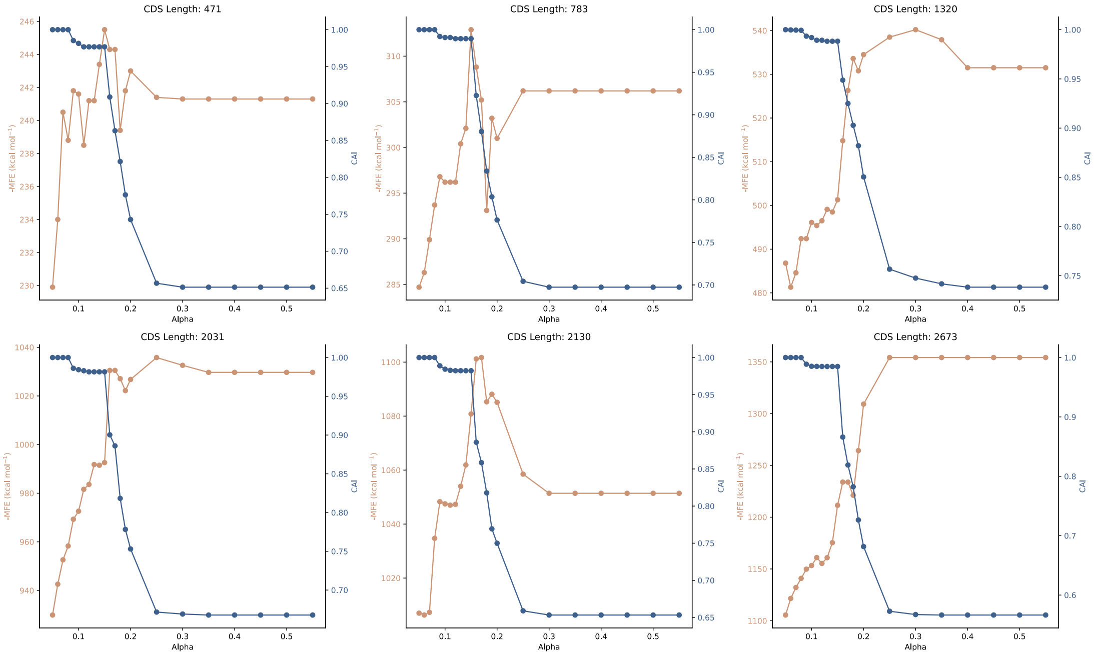


**Supplementary Figure 2.** The influence of MFE-CAI weight alpha on MFE and CAI.

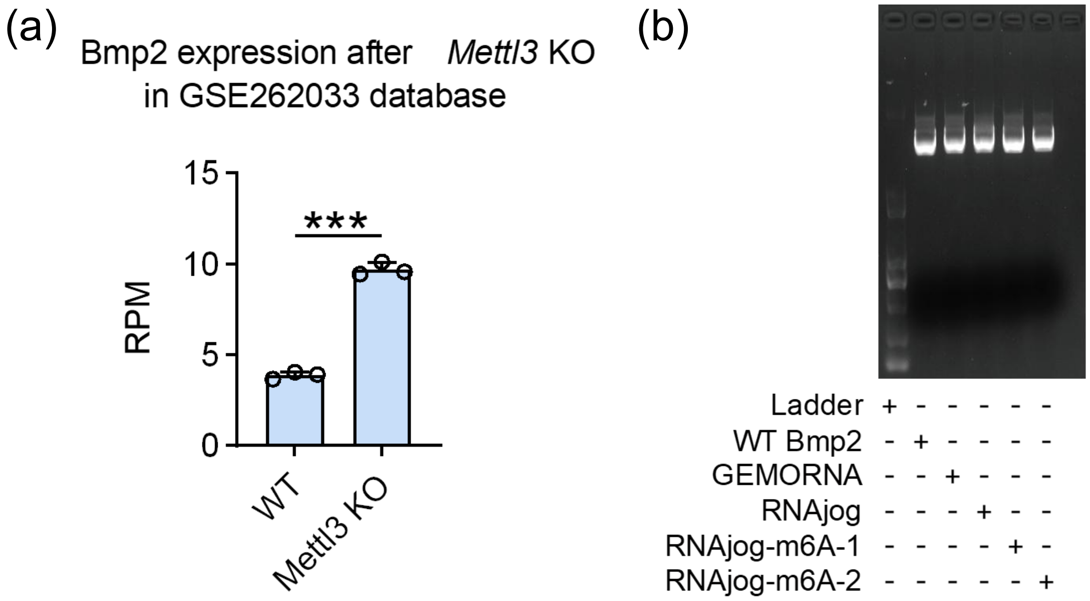


**Supplementary Figure 3.** (a)The RPM (Reads per Million) of Bmp2 after Mettl3 knockout. *P*-values were calculated by Student's t-test. (*** $p < 0.001$, $n = 3$). (b) Validation of plasmid loading for transfection. Agarose gel electrophoresis was performed to verify the integrity and quantity of the optimized plasmids. Consistent loading was ensured by combining spectrophotometric concentration measurements with visual confirmation of band intensity on the gel, ensuring that equal masses of plasmids were used for each transfection group.

#### Supplementary Table 1

The Comparison of functional features across different RNA design models.

| Model | Top-k output[1] | Controllable Weight[2] | Dataset-Free[3] | Extensibility[4] | Exclude RE site[5] | Exclude m6a motif[6] |
|---|---|---|---|---|---|---|
| RiboCode | ✓ | ✓ | | | | |
| CodonTransformer | ✓ | | | | | |
| LinearDesign | | ✓ | ✓ | | ✓[7] | |
| GEMORNA | ✓ | | | | | |
| RNAJog | ✓ | ✓ | | ✓ | ✓ | ✓ |
| RNAJog-Zero | ✓ | ✓ | ✓ | ✓ | ✓ | ✓ |

[1] Ability to generate top-k optimal solutions.

[2] Controllable multi-objective weighting.

[3] Ability to optimize without training data.

[4] Whether new optimization objectives can be easily added.

[5] Ability to optimize codons while excluding restriction enzyme cutting sites.

[6] Ability to optimize codons while excluding m6a motifs.

[7] As of the time of writing, this feature of LinearDesign has not yet been open-sourced.

#### Supplementary Table 2

The selected six representative mRNA sequences:

| Length | Sequence |
|---|---|
| 471 | AUGCUCGGGUCUCCUUGCGGCCCCCAGCUCAGCGACCGAGACGCAGACGAGGACCAGUG<br>UUCACGCGAGUUCAGGGGGCGGCGUAGCCGCCGCCCGCCCAGGAGGACCAUGUUGCGC<br>GGCAAGUCCCGGCUCAACGUGGAGUGGCUGGGCUACUCGCCAGGCCUGCUCCUCGAGC<br>ACAGGCCCCUCCUGGCAGGGCGCACGCCGCGGAGCCACCGCCGAAAUGAAAGCUCAUUA<br>GCAUCUACACUGAAGACGCUCCUGUUCUUCACAGCUUUAAUGAUCACUGUUCCUAUUG<br>GGUUAUAUUUCACAACUAAAUCUUACAUAUUUGAAGGCGCCCUUGGGAUGUCCAAUAG<br>GGACAGCUAUUUUUACGCUGCUAUUGUUGCAGUGGUCGCCGUCCAUGUGGUGCUGGCC<br>CUCUUUGUGUAUGUGGCCUGGAAUGAAGGCUCACGACAGUGGCGUGAAGGCAAACAGG<br>AUUAA |
| 783 | AUGGCGAUUCUUUUUGCUGUUGUUGCCAGGGGGACCACUAUCCUUGCCAAACAUGCUU<br>GGUGUGGAGGAAACUUCCUGGAGGUGACAGAGCAGAUUCUGGCUAAGAUACCUUCUGA<br>AAAUAACAAACUAACGUACUCACAUGGCAAUUAUUUGUUUCAUUACAUCUGCCAAGACA<br>GGAUUGUAUAUCUUUGUAUCACUGAUGAUGAUUUUGAACGUUCCCGAGCCUUUAAUUU<br>UCUGAAUGAGAUAAAGAAGAGGUUCCAGACUACUUACGGUUCAAGAGCACAGACAGCAC<br>UUCCAUAUGCCAUGAAUAGCGAGUUCUCAAGUGUCUUAGCUGCACAGCUGAAGCAUCA<br>CUCUGAGAAUAAGGGCCUAGACAAAGUGAUGGAGACUCAAGCCCAAGUGGAUGAACUG<br>AAAGGAAUCAUGGUCAGAAACAUAGUCUGUCACCUUCAAAACUACCAGCAGAAAUCUUG<br>CUCGAGCCAUGUGUAUGAAGAACCUCAAGCUCACUAUUAUCAUCAUCAUCGUAUCAAUU<br>GUGUUCAUCUAUAUCAUUGUUUCACCUCUCUGUGGUGGAUUUACAUGGCCAAGCUGUG<br>UGAAGAAAUAGGAAAGAAGAAGUUACCAUUAACCAAGGAUAUGAGAGAACAAGGAGUU<br>AAAAGCAAUCCAUGUGACUCAAGCCUUUCACAUACUGACAGAUGGUAUCUGCCAGUCUC<br>UUCAACCCUCUUCUCACUUUUUAAAAUCUUGUUCCAUGCCUCCAGGUUUAUCUUUGUC<br>UUAUCUACCAGUUUAUUCCUGUGA |
| 1320 | AUGGAGGAGAGGAAACCAGCUCAUGUGCUGAGAAGUUUUAAAUAUGCUGCAUUUAUGA<br>AUGAAGAGCUUCGAAACUUGUCUUUGUCUGGCCAUGUGGGAUUUGACAGCCUCCCUGA<br>CCAGCUGGUCAACAAGUCUACUUCUCAAGGAUUCUGUUUCAACAUCCUUUGUGUUGGU<br>GAGACAGGCAUUGGCAAAUCCACGUUAAUGGACACUUUGUUCAACACCAAAUUUGAAA<br>GUGACCCAGCUACUCACAAUGAACCAGGUGUUCGGUUAAAAGCCAGAAGUUAUGAGCU<br>UCAGGAAAGCAAUGUACGGCUGAAGUUAACCAUUGUUGACACCGUGGGAUUUGGAGAC<br>CAGAUAAAUAAAGAUGACAGCUAUAAGCCGAUAGUAGAAUAUAUUGAUGCCCAGUUCG<br>AGGCCUACCUGCAAGAGGAAUUGAAGAUUAAACGUUCUCUCUUCAACUACCAUGACACG<br>AGGAUCCAUGCCUGCCUCUACUUUAUUGCCCCUACUGGACAUUCACUAAAGUCCCUGGA<br>UCUGGUCACCAUGAAAAAGCUGGACAGUAAGGUGAACAUCAUUCCAAUAAUUGCAAAAG<br>CUGACACCAUUGCCAAGAAUGAACUGCACAAAUUCAAGAGUAAGAUCAUGAGUGAACUG<br>GUCAGCAAUGGGGUCCAGAUAUAUCAGUUUCCCACUGAUGAAGAAACGGUGGCAGAGA<br>UUAACGCAACAAUGAGUGUCCAUCUCCCAUUUGCAGUGGUUGGCAGCACCGAAGAGGU<br>GAAGAUUGGCAACAAGAUGGCAAAGGCCAGGCAGUACCCCUGGGGUGUGGUGCAGGUU<br>GAGAAUGAAAAUCAUUGCGAUUUUGUGAAACUUCGAGAGAUGCUGAUCCGCGUGAACA<br>UGGAGGACUUGCGAGAGCAGACUCACACCCGCCACUAUGAAUUGUACCGACGCUGUAA |

| | |
|---|---|
| | GCUUGAAGAGAUGGGGUUCAAGGACACUGACCCUGACAGCAAACCCUUCAGUCUUCAG<br>GAGACAUAUGAAGCAAAAAGGAAUGAAUUCCUGGGAGAACUGCAGAAGAAAGAAGAAG<br>AAAUGAGACAAAUGUUUGUUAUGAGAGUGAAGGAGAAAGAAGCUGAACUUAAGGAGGC<br>AGAGAAAGAGCUUCACGAGAAGUUUGACCUUCUAAAGCGGACACACCAAGAAGAAAAGA<br>AGAAAGUGGAAGACAAGAAGAAGGAGCUUGAGGAGGAGGUGAACAACUUCCAGAAGAA<br>GAAAGCAGCGGCUCAGUUACUACAGUCCCAGGCCCAGCAAUCUGGGGCCCAGCAAACCA<br>AGAAAGACAAGGAUAAGAAAAAUGCAAGCUUCACAUAA |
| 2031 | AUGUCGCGCCGGGCCCUCCGGAGGCUGAGGGGGAACAGCGCGGCCAGGAGCCCCUCG<br>GGCCCGGCGCCUUGCAUUUCGAUCUCCGUGAUGACGAUGACGCGGAAGAAGAAGGGCC<br>CAAGCGGGAGCUUGGUGUCCGGCGUCCCGGGGGCGCAGGGAAGGAGGGCGUCCGAGUC<br>AACAACCGCUUCGAGCUGAUAAACAUUGACGAUCUUGAGGAUGACCCUGUGGUGAACG<br>GGGAGAGGUCUGGCUGUGCGCUCACAGACGCUGUGGCACCAGGGAACAAAGGAAGGGG<br>UCAGCGUGGAAACACAGAGAGCAAGACGGAUGGAGAUGACACCGAGACAGUGCCCUCA<br>GAGCAGUCUCAUGCAAGUGGCAAACUCCGGAAGAAGAAAAAAAAACAGAAAAACAAGAA<br>AAGCAGCACGGGAGAAGCAUCGGAAAACGGACUAGAAGAUAUCGAUCGCAUCCUAGAGA<br>GGAUUGAGGACAGCACUGGGUUGAACCGUCCCGGCCCAGCUCCCCUGAGCUCCAGGAA<br>GCACGUUCUCUACGUGGAGCACAGACACUUGAAUCCAGACACAGAACUGAAAAGGUAUU<br>UUGGUGCCCGGGCAAUCCUGGGGGAGCAAAGGCCACGGCAGAGACAACGUGUGUACCC<br>CAAGUGCACAUGGCUGACCACCCCUAAAAGCACCUGGCCCCGCUACAGCAAACCAGGUC<br>UGUCCAUGCGGCUGCUGGAAUCAAAAAAAGGCCUCUCCUUCUUUGCGUUUGAGCACAG<br>UGAGGAGUACCAGCAGGCUCAGCACAAGUUCCUGGUGGCCGUGGAGUCUAUGGAGCCG<br>AACAACAUCGUGGUUCUGCUCCAGACGAGCCCUUACCACGUUGACUCACUCCUGCAGCU<br>CAGCGAUGCCUGCCGCUUUCAAGAGGAUCAGGAGAUGGCUCGAGACCUCGUAGAGAGA<br>GCGCUGUACAGCAUGGAAUGUGCGUUCCACCCCCUGUUCAGUCUCACCAGUGGGGCCU<br>GCCGGCUGGAUUACCGCAGACCCGAGAACAGGAGCUUCUACCUGGCCCUCUACAAGCAG<br>AUGAGCUUCCUGGAGAAGCGAGGCUGCCCGCGCACGGCGCUGGAGUACUGCAAGCUCA<br>UCCUGAGUCUCGAGCCGGAUGAGGACCCCCUCUGCAUGCUGCUGCUCAUCGACCACCUG<br>GCCUUGCGGGCCCGGAACUACGAGUACCUGAUCCGCCUCUUCCAGGAGUGGGAGGCUC<br>AUCGGAACCUGUCCCAGCUCCCUAAUUUUGCCUUCUCUGUUCCACUGGCGUAUUUCCU<br>GCUGAGCCAGCAGACAGACCUCCCUGAGUGUGAGCAGAGCUCUGCCAGGCAGAAGGCC<br>UCUCUCCUGAUACAGCAGGCGCUCACCAUGUUCCCUGGAGUCCUCCUGCCCCUGCUCGA<br>GUCUUGCAGUGUGCGGCCCGACGCCAGCGUUUCCAGUCACCGCUUCUUUGGACCCAAU<br>GCUGAAAUAAGCCAGCCCCCUGCCCUGAGCCAGCUGGUGAACCUGUACCUUGGGAGGUC<br>ACACUUUCUCUGGAAAGAGCCCGCCACCAUGAGCUGGCUGGAGGAGAACGUCCACGAG<br>GUUCUGCAAGCAGUGGACGCCGGGGACCCAGCCGUGGAAGCCUGUGAGAACCGGCGGA<br>AGGUGCUCUACCAGCGUGCACCCAGGAAUAUCCACCGCCAUGUGAUCCUCUCUGAGAUC<br>AAGGAAGCCGUCGCUGCCCUGCCCCCGGACGUGACCACGCAGUCUGUGAUGGGGUUUG<br>AUCCUCUGCCUCCUUCGGACACAAUCUACUCCUACGUCAGGCCAGAGAGGCUAAGUCCU<br>AUCAGCCAUGGAAACACCAUUGCUCUCUUCUUCCGGUCACUGUUGCCAAACUAUACCAU<br>GGAGGGGGAGAGGCCCGAGGAAGGAGUGGCUGGGGGUCUGAACCGCAACCAGGGCCUG<br>AACAGGCUGAUGCUGGCUGUGCGCGACAUGAUGGCCAACUUCCACCUCAACGACCUGGA<br>GGCGCCGCACGAGGACGACGCUGAGGGGGAGGGGGAGUGGGACUGA |
| 2130 | AUGGCGGCCAACAUGUACCGGGUCGGAGAUUAUGUCUUCUUUGAGAAUUCUUCAAGCA<br>ACCCCUAUCUGAUCAGGCGAAUAGAGGAACUGAAUAAGACUGCAAGCGGAAAUGUGGA |

| | |
|---|---|
| | GGCAAAGGUGGUUUGUUUCUACAGGAGACGGGACAUCUCUCAGAGCCUCAUCCAGCUG<br>GCAGACAAACACGCAAAGGACCUGGAAGAAGAGAAGGAAAGCCCCCCAGAGUCCGACCU<br>CACUGAAAAACAGAAACAUCAGCUCCGCCACAGAGAGCUUUUCCUCUCCCGCCAGUACG<br>AAUCCCUCCCUGCCACGCACAUCAGGGGGAAGUGCAGCGUGGCCCUCCUUAACGAGACA<br>GAAGUUGUGCUCUCCUACCUAGAGAAAGAGGACACAUUUUUUACUCAUUGGUGUACG<br>ACCCAACACAGAAGACGCUAUUGGCGGACAAGGGGGAGAUUCGCGUUGGUCCACGUUU<br>UCAGGCUGAUGUCCCUGAAAUGCUACAGGAAGGAGAAGCUGAUGACAGAGACCAGUCC<br>AAACUGGAGAUGAAGAUGUGGGAUCCAGAGUGUCCGCUGACCAACAAACAGAUCGACCA<br>GUUUCUUGUGGUUGCCCGAGCGGUUGGUACGUUUGCCCGGGCGUUGGACUGCAGCAGC<br>UCCGUCAGACAGCCCAGCUUACACAUGAGCGCCGCGGCAGCCUCACGAGACAUCACACU<br>GUUUCACGCUAUGGACACACUGCAUCGGCAUGGCUAUGACCUGUCCAGUGCUCUCAGC<br>GUUCUUGUUCCUCAAGGUGGGCCAGUCCUGUGCCGGGAUGAGAUGGAGGAGUGGAGCU<br>CGUCAGAGGCCAAUCUGUUUGAAGAGGCCCUGGAGAAAUACGGCAAAGACUUCAACGA<br>UAUACGGCAAGAUUUUCUUCCCUGGAAGUCGCUGACCAGCAUCAUUGAGUAUUAUUAC<br>AUGUGGAAAACCACAGACAGAUACGUUCAGCAGAAACGACUGAAGGCAGCUGAAGCAGA<br>GAGCAAACUGAAGCAAGUCUACAUCCCUACAUACAAUAAACCCAACCCAAACCAGAUCU<br>CUGUCAGCAAUGGCAAGAUGGCAACAGUGAACGGUGCAGCAGCAGGCACAGGCAGUUU<br>CCACACAGCUGGAGGGGGCCGCGCAUGCGAGAGCUGUUUUGCUGUACAGUCUGCUCAG<br>UGGUACUCCUGGGGUCCUCCUAAUAUGCAGUGUCGCCUGUGUGUUUCCUGUUGGAUGU<br>ACUGGAAGAAGUACGGUGGCUUGAAGAUGCCAAGCAGAGCUGAGGGGGCAGAGGAAAA<br>AACACCUCCAAGCCCUGCACCUAACGAAUCACGUUCUCGUGGCCACUGUGCCCGUCAGU<br>CUUCCCACAUGGUCCCCAUCCGGAACAGUGGUAGCCCAAAAUCCUCCAUGAAGACCAAG<br>CAGGCGUUCCUCCUGCAGGCAACGCGCCUCACCAAGCUGGCGCGUCACAUGUGCCGCGA<br>UCUCAUCAGGCUGCGCCGGGCCGCGCGCCGCCCCUUUGUGCCCAUUAACUGUGGUGCCA<br>UAAAGGCGGAGUAUAUGAUCCGGGUGUCCGAGGGCAUGACAGGGAGACCUUUGAAGCC<br>CAAAUCUUCUCCCAGGAGCACCCUGACCAGCGUCUUGCAGUAUCUUGAGUCCCGUCCAG<br>CAACUCAUGUUCAACGGCCUCAUCGCACCCCUGGCCUACAGGUGCAGCCUCCACGACGC<br>CUGCUUUCCUCUCUUCCAAGUCAUGGCCCGCUUGGAAUGUUGGGAAAACGAAGCUACC<br>AUCACCACAGCAGGGUUGAAUCUGCAGAAAGGAGGGCUGGUGCUACAGGACAAGAAAA<br>UCCAGCUCAUAUUGUGGGGCCCAUUCUGCAGCAUAAUGGGAGCAGCACCGGUGGCUCC<br>AGCCUUCGUGGCAGUGGCUUGAUGCUCCGUAAGAGACGACCCAACUGGAUCGAUGCUC<br>CUGAUGACAGCUUCUUCCUGGUUUCCCGAGAGACGAGAAAAGCCAGGAGGAUGCUGUC<br>CCGUUCACAGUUGAGAAGAGCAUGUAGACAACCAUGUGAGCAGAUCAGUCUGCGCAGG<br>GUCCCCCAGGGGCCCUCACAGGUACCCAUCCUGGCACCUCCUCACCCCAGUUUACGGAU<br>GCGCGGCCCAAUCGUCAUCCAUGACUGA |
| 2673 | AUGGGGCUCACCACGGGUCCAAAUGGAGCUCGUGCUGCCCCACAGAAAAGUUUGCGUC<br>GAGUGUGGCAGCGGGGCAGUCGUGGUUCAAAGGUGAAGCAUCGGGGCGGCACAAUGGA<br>GGGUGAGCAGAGCACUCCUGCCGCCAGUGCCAGCAGCACUACGGGUACCAGCACAUCCA<br>CUACAAACUCCACAGGCACCACCAGCGCCUCCACUGCCAGCACUGCCAGCUCCACCAGCA<br>GCAGCACGAGCAGCGCGAGACAGGCCGUUCCUCAGAUAUCUGUAUACAGCGGCAUACCU<br>GACCGACAGACCGUACAGGUGAUCCAGCAGGCUCUGAAUAGGCAGCCCAGCACCGCAGC<br>GCAGUACCUGCAGCAGAUGUAUGCAGCUCAACAGCAGCAUCUGAUGUUGCAAACAGCAG<br>CCCUGCAGCAGCAGCACCUCAGCACAGCACAGCUUCAGAGCCUCGCAGCAGUCCAGCAG<br>GCUAGUUUAGCAGCCGGUAGACAGAGUUCAAACCAGAAUGGGACCUCAUCUACUCAGA |

| | |
|---|---|
| | GUGGAUCAUCACAGACGACAAUUAAUUUAACCACAUCUCCUGCUGCCGCUCAGCUGAUC<br>AGUCGUGCCCAAAGUGUAAACUCCGCCCCAUCUGGGAUUUCCCAGCAAGCGGUGUUGC<br>UUGGCAACGCCUCGAGCUCCACCCUGACAGCCAGCCAAGCUCAAAUGUACCUGCGGGCA<br>CAGAUGGCUCAGCAAAGUAAUCUUGUGCAUGUAGCCAGAAGUCUGGGUCGAGCCGUCC<br>CUCUGUCUUCUCAGCUGAUCCUGGCCCCGACUGCUACAGUAACUGCUCUACAGUCUGAC<br>ACUAACACAACCAAUAACACACAGUCUGCCUCUGCACAGGUGCAGAACCUUGCCCUGCG<br>CAACCACCAGGGGGUGUCUUCCACCCCUCAGGCCACAGGACUCCAGGCACUGAGUCUAA<br>AGCAGACCCCUGUCUCCAUUCAGCCCACCCCACUUAUCAAGACACCCAGCCAGAGCACU<br>GGUGCCGGGAAAAACAACCUGAACGCUGCUUCCUCGGAUGGAGUGAAGAAGGCAGAGG<br>AGGGACAAACCGAGAUGAAAGCAGUGAACAUGAGUCGUAAUGUGUCCUCUUCACAUCCC<br>UUAAUAGCACCAGCCUAUGCUCAAAUCCAGCCCCAGGCUCUGAUUAAGCAACAACCUCA<br>ACAGUUUGUCAUUCAACCUCAAGCCACUCCCACCUCCAGAAGCCAACCUCAGUUACUAC<br>AGACCGCCUCAAUCCAGCCCCACCAACAAACAGCCUCGCUGGGUAUCCCAAGCUCCGCC<br>CAGACCUCAACCGUAACGGGCAUCAUUCCAGUUUUACCCAAACCAGCUCUUCACAACCA<br>GGGAUCAGCUCAAACGCAACAGGCCACCAUCUUCCAUUCAACCGUAAGCCACCAUGCAC<br>ACACAGCUCUGGCCCAUGCUAAAGCUCAGCCGGUCCAACUGACAGCCAUCAAUCUCCAA<br>AUACAACCAGCACAGAACCAGGCUUCAAGGCUUGCUCAGGAUGACAAGGAGAAGCCCAC<br>CUCUUUGGUCUUAAGAGAGAUUUGCCCACCAGUACAAAUCCAUCCAUCUACUACAACCC<br>AAGGCACCGACCCACCUGAACAAUCCAAAGCUGACACUGUUAAUGCUACAUCUCAAACA<br>CAAGGCUCAGUGGAAACGGCGCGGGAGCCCACGGGAGCACAGCUGACAUCCAGCAUGAC<br>CUCCGCAGUCUCCAGUGAGCCUCCGCCGGCAGCUGUGAUGGGAAAUACCACACAAAACG<br>GAGAGAACAAACCACCUCAAGCCAUAGUGAAACCACAUGUUCUGACGCAUGUCAUUGAG<br>GGCUUUGUUAUCCAGGAGGGAGCCGAGCCAUUCCCUGUGGAAAGACCUCCGGUGGCGG<br>CCGAAAGCCCCAAAAAACCAGACACGCAGCUGCCCUCAGAUCCAGAGAAAACGGCCAGC<br>AGCAACUCCCUCAGUGCCACAAACUCAGACACAGAGGAGGCUGGACAACAAGACACAAG<br>AGUUGAGGAGGAGACAGAGAAGCCCAAGCUCACAUGUGAGCUCUGUGGAUGGGUGGAU<br>UUCGCUCACAAGUUUAAACGCUCAAAGCGUUUCUGUUCCAUGGUCUGUGCUAAAAGGU<br>ACAGUGUAAGUUGCACGAAGCGGGUUGGUCUGUUCCGCCCAGACAGAACUAAAUCGAC<br>UAAUCAGAUAAAUCAGUGGAGGAGGAGGAGGUCUCAUAGUCCCAGAGGGAGGGAGGCC<br>AAAAAGCAGCGGCUUUCUGUAUCUCAGCAAUCUCAGGGAGAAUCUAUAUCAUCACCUCU<br>UCAGUCUCAGCCUAGUCAGGGAGAGUCAAGUCCGUGUUCGGAUAUAUCCAGUUAUGAG<br>GAAGCCACUUCCCCUCUCUCUGCUGCCAGCUCUGGGCCAGUGGUUUCUCAAGGUGUGG<br>UUGCUGCUAAUGAACCAGAAGAGCUUCCUCUCCUCAGUCAGCACUUCCUGCCCUGUGAC<br>CCCACUAAAUGGAAUGUGCAAGAAGUGUUUGAGUUCAUCCGCUCGUUGCCAGGUUGCC<br>AGGAGAUUGCAGAUGAGUUUCGUGCUCAGGAGAUUGAUGGACAGGCGCUGCUGCUACU<br>GAAAGAAGACCAUCUAAUGAGUGCCAUGAAUAUUAAACUGGGACCCGCCCUCAAGAUAU<br>UCGCUCGCAUAAACAUGCUGAAAGAUUCCUAG |

#### Supplementary Table 3

| | length | alpha | MFE | CAI |
|---|---|---|---|---|
| RNAJog optimized mRNA | 4107 | 0.11 | -1678.6 | 0.983254 |
| Reference mRNA | 4107 | / | -963.5 | 0.639461 |

| | Sequence |
|---|---|
| RNAJog optimized mRNA | AUGGACAAGAAGUACUCCAUCGGGCUGGACAUCGGGACCAACAGCGUGGGGUGGGCCGUGAUCACCGACGAGUACAAGGUGCCCAGCAAGAAGUUCAAGGUGCUGG<br>GGAACACCGACCGGCACAGCAUCAAGAAGAACCUGAUCGGGGCCCUGCUGUUCGACAGCGGGGAGACCGCCGAGGCCACCCGGCUGAAGCGGACCGCCCGGCGGCGG<br>UACACCCGGCGGAAGAACCGGAUCUGCUACCUGCAGGAGAUCUUCAGCAACGAGAUGGCCAAGGUGGACGACAGCUUCUUCCACCGGCUGGAGGAGUCCUUCCUGGU<br>GGAGGAGGACAAGAAGCACGAGCGGCACCCCAUCUUCGGGAACAUCGUGGACGAGGUGGCCUACCACGAGAAGUACCCCACCAUCUACCACCUGCGGAAGAAGCUGG<br>UGGACAGCACCGACAAGGCCGACCUGCGGCUGAUCUACCUGGCCCUGGCCCACAUGAUCAAGUUCCGGGGGCACUUCCUGAUCGAGGGGGACCUGAACCCCGACAAC<br>AGCGACGUGGACAAGCUGUUCAUCCAGCUGGUGCAGACCUACAACCAGCUGUUCGAGGAGAACCCCAUCAACGCCAGCGGGGUGGACGCCAAGGCCAUCCUGAGCGC<br>CCGGCUGAGCAAGAGCCGGCGGCUGGAGAACCUGAUCGCCCAGCUGCCCGGGGAGAAGAAGAACGGGCUGUUCGGGAACCUGAUCGCCCUGAGCCUGGGGCUGACC<br>CCCAACUUCAAGAGCAACUUCGACCUGGCCGAGGACGCCAAGCUGCAGCUGAGCAAGGACACCUACGACGACGACCUGGACAACCUGCUGGCCCAGAUCGGGGACCA<br>GUACGCCGACCUGUUCCUGGCCGCCAAGAACCUGUCCGACGCCAUCCUGCUGAGCGACAUCCUGCGGGUGAACACCGAGAUCACCAAGGCCCCCCUGAGCGCCAGCA<br>UGAUCAAGCGGUACGACGAGCACCACCAGGACCUGACCCUGCUGAAGGCCCUGGUGCGGCAGCAGCUGCCCGAGAAGUACAAGGAGAUCUUCUUCGACCAGAGCAAG<br>AACGGGUACGCCGGGUACAUCGACGGGGGGGCCAGCCAGGAGGAGUUCUACAAGUUCAUCAAGCCCAUCCUGGAGAAGAUGGACGGGACCGAGGAGCUGCUGGUGA<br>AGCUGAACCGGGAGGACCUGCUGCGGAAGCAGCGGACCUUCGACAACGGGAGCAUCCCCCACCAGAUCCACCUGGGGGAGCUGCACGCCAUCCUGCGGCGGCAGGAG<br>GACUUCUACCCCUUCCUGAAGGACAACCGGGAGAAGAUCGAGAAGAUCCUGACCUUCCGGAUCCCCUACUACGUGGGGCCCCUGGCCCGGGGGAACAGCCGGUUCGC<br>CUGGAUGACCCGGAAGUCCGAGGAGACCAUCACCCCCUGGAACUUCGAGGAGGUGGUGGACAAGGGGGCCAGCGCCCAGAGCUUCAUCGAGCGGAUGACCAACUUCG<br>ACAAGAACCUGCCCAACGAGAAGGUGCUGCCCAAGCACAGCCUGCUGUACGAGUACUUCACCGUGUACAACGAGCUGACCAAGGUGAAGUACGUGACCGAGGGCAUG<br>CGGAAGCCCGCCUUCCUGUCCGGGGAGCAGAAGAAGGCCAUCGUGGACCUGCUGUUCAAGACCAACCGGAAGGUGACCGUGAAGCAGCUGAAGGAGGACUACUUCAA<br>GAAGAUCGAGUGCUUCGACAGCGUGGAGAUCAGCGGGGUGGAGGACCGGUUCAACGCCAGCCUGGGGACCUACCACGACCUGCUGAAGAUCAUCAAGGACAAGGAC<br>UUCCUGGACAACGAGGAGAACGAGGACAUCCUGGAGGACAUCGUGCUGACCCUGACCCUGUUCGAGGACCGGGAGAUGAUCGAGGAGCGGCUGAAGACCUACGCCC<br>ACCUGUUCGACGACAAGGUGAUGAAGCAGCUGAAGCGGCGGCGGUACACCGGGUGGGGGCGGCUGUCCCGGAAGCUGAUCAACGGGAUCCGGGACAAGCAGAGCGG<br>GAAGACCAUCCUGGACUUCCUGAAGUCGGACGGGUUCGCCAACCGGAACUUCAUGCAGCUGAUCCACGACGACUCCCUGACCUUCAAGGAGGACAUCCAGAAGGCCC<br>AGGUGAGCGGGCAGGGGGACAGCCUGCACGAGCACAUCGCCAACCUGGCCGGCAGCCCCGCCAUCAAGAAGGGGAUCCUGCAGACCGUGAAGGUGGUGGACGAGCU<br>GGUGAAGGUGAUGGGGCGGCACAAGCCCGAGAACAUCGUGAUCGAGAUGGCCCGGGAGAACCAGACCACCCAGAAGGGGCAGAAGAACUCCCGGGAGCGGAUGAAG<br>CGGAUCGAGGAGGGGAUCAAGGAGCUGGGGAGCCAGAUCCUGAAGGAGCACCCCGUGGAGAACACCCAGCUGCAGAACGAGAAGCUGUACCUGUACUACCUGCAGAA<br>CGGGCGGGACAUGUACGUGGACCAGGAGCUGGACAUCAACCGGCUGAGCGACUACGACGUGGACCACAUCGUGCCCCAGAGCUUCCUGAAGGACGACAGCAUCGACA<br>ACAAGGUGCUGACCCGGAGCGACAAGAACCGGGGGAAGUCCGACAACGUGCCCAGCGAGGAGGUGGUGAAGAAGAUGAAGAACUACUGGCGGCAGCUGCUGAACGC<br>CAAGCUGAUCACCCAGCGGAAGUUCGACAACCUGACCAAGGCCGAGCGGGGGGGGCUGAGCGAGCUGGACAAGGCCGGGUUCAUCAAGCGGCAGCUGGUGGAGACC<br>CGGCAGAUCACCAAGCACGUGGCCCAGAUCCUGGACAGCCGGAUGAACACCAAGUACGACGAGAACGACAAGCUGAUCCGGGAGGUGAAGGUGAUCACCCUGAAGAG<br>CAAGCUGGUGAGCGACUUCCGGAAGGACUUCCAGUUCUACAAGGUGCGGGAGAUCAACAACUACCACCACGCCCACGACGCCUACCUGAACGCCGUGGUGGGGACCG<br>CCCUGAUCAAGAAGUACCCCAAGCUGGAGAGCGAGUUCGUGUACGGGGACUACAAGGUGUACGACGUGCGGAAGAUGAUCGCCAAGAGCGAGCAGGAGAUCGGGAA<br>GGCCACCGCCAAGUACUUCUUCUACAGCAACAUCAUGACUUCUUCAAGACCGAGAUCACCCUGGCCAACGGGGAGAUCCGGAAGCGGCCCCUGAUCGAGACCAACGG<br>GGAGACCGGGGAGAUCGUGUGGGACAAGGGGCGGGACUUCGCCACCGUGCGGAAGGUGCUGAGCAUGCCCCAGGUGAACAUCGUGAAGAAGACCGAGGUGCAGACC<br>GGGGGGUUCAGCAAGGAGAGCAUCCUGCCCAAGCGGAACAGCGACAAGCUGAUCGCCCGGAAGAAGGACUGGGACCCCAAGAAGUACGGGGGGUUCGACAGCCCCAC<br>CGUGGCCUACAGCGUGCUGGUGGUGGCCAAGGUGGAGAAGGGGAAGAGCAAGAAGCUGAAGAGCGUGAAGGAGCUGCUGGGGAUCACCAUCAUGGAGCGGAGCAGC<br>UUCGAGAAGAACCCCAUCGACUUCCUGGAGGCCAAGGGGUACAAGGAGGUGAAGAAGGACCUGAUCAUCAAGCUGCCCAAGUACUCCCUGUUCGAGCUGGAGAACGG<br>GCGGAAGCGGAUGCUGGCCUCCGCCGGGGAGCUGCAGAAGGGGAACGAGCUGGCCCUGCCCAGCAAGUACGUGAACUUCCUGUACCUGGCCAGCCACUACGAGAAGC<br>UGAAGGGGAGCCCCGAGGACAACGAGCAGAAGCAGCUGUUCGUGGAGCAGCACAAGCACUACCUGGACGAGAUCAUCGAGCAGAUCAGCGAGUUCUCCAAGCGGGU<br>GAUCCUGGCCGACGCCAACCUGGACAAGGUGCUGUCCGCCUACAACAAGCACCGGGACAAGCCCAUCCGGGAGCAGGCCGAGAACAUCAUCCACCUGUUCACCCUGA |

| | |
|---|---|
| | CCAACCUGGGGGCCCCCGCCGCCUUCAAGUACUUCGACACCACCAUCGACCGGAAGCGGUACACCAGCACCAAGGAGGUGCUGGACGCCACCCUGAUCCACCAGAGCA<br>UCACCGGGCUGUACGAGACCCGGAUCGACCUGAGCCAGCUGGGGGGGGACUGA |
| Reference mRNA | AUGGAUAAGAAAUACUCAAUAGGCUUAGAUAUCGGCACAAAUAGCGUCGGAUGGGCGGUGAUCACUGAUGAAUAUAAGGUUCCGUCUAAAAAGUUCAAGGUUCUGG<br>GAAAUACAGACCGCCACAGUAUCAAAAAAAAUCUUAUAGGGGCUCUUUUAUUUGACAGUGGAGAGACAGCGGAAGCGACUCGUCUCAAACGGACAGCUCGUAGAAGG<br>UAUACACGUCGGAAGAAUCGUAUUUGUUAUCUACAGGAGAUUUUUUCAAAUGAGAUGGCGAAAGUAGAUGAUAGUUUCUUUCAUCGACUUGAAGAGUCUUUUUUGG<br>UGGAAGAAGACAAGAAGCAUGAACGUCAUCCUAUUUUUGGAAAUAUAGUAGAUGAAGUUGCUUAUCAUGAGAAAUAUCCAACUAUCUAUCAUCUGCGAAAAAAAUU<br>GGUAGAUUCUACUGAUAAAGCGGAUUUGCGCUUAAUCUAUUUGGCCUUAGCGCAUAUGAUUAAGUUUCGUGGUCAUUUUUUGAUUGAGGGAGAUUUAAAUCCUGAU<br>AAUAGUGAUGUGGACAAACUAUUUAUCCAGUUGGUACAAACCUACAAUCAAUUAUUUGAAGAAAACCCUAUUAACGCAAGUGGAGUAGAUGCUAAAGCGAUUCUUU<br>CUGCACGAUUGAGUAAAUCAAGACGAUUAGAAAAUCUCAUUGCUCAGCUCCCCGGUGAGAAGAAAAAUGGCUUAUUUGGGAAUCUCAUUGCUUUGUCAUUGGGUUU<br>GACCCCUAAUUUUAAAUCAAAUUUUGAUUUGGCAGAAGAUGCUAAAUUACAGCUUUCAAAAGAUACUUACGAUGAUGAUUUAGAUAAUUUAUUGGCGCAAAUUGGA<br>GAUCAAUAUGCUGAUUUGUUUUUGGCAGCUAAGAAUUUAUCAGAUGCUAUUUUACUUUCAGAUAUCCUAAGAGUAAAUACUGAAAUAACUAAGGCUCCCCUAUCAG<br>CUUCAAUGAUUAAACGCUACGAUGAACAUCAUCAAGACUUGACUCUUUUAAAAGCUUUAGUUCGACAACAACUUCCAGAAAAGUAUAAAGAAAUCUUUUUUGAUCAA<br>UCAAAAAACGGAUAUGCAGGUUAUAUUGAUGGGGGAGCUAGCCAAGAAGAAUUUUAUAAAUUUAUCAAACCAAUUUUAGAAAAAAUGGAUGGUACUGAGGAAUUAU<br>UGGUGAAACUAAAUCGUGAAGAUUUGCUGCGCAAGCAACGGACCUUUGACAACGGCUCUAUUCCCCAUCAAAUUCACUUGGGUGAGCUGCAUGCUAUUUUGAGAAG<br>ACAAGAAGACUUUUAUCCAUUUUUAAAAGACAAUCGUGAGAAGAUUGAAAAAAUCUUGACUUUUCGAAUUCCUUAUUAUGUUGGUCCAUUGGCGCGUGGCAAUAGU<br>CGUUUUGCAUGGAUGACUCGGAAGUCUGAAGAAACAAUUACCCCAUGGAAUUUUGAAGAAGUUGUCGAUAAAGGUGCUUCAGCUCAAUCAUUUAUUGAACGCAUGA<br>CAAACUUUGAUAAAAAUCUUCCAAAUGAAAAAGUACUACCAAAACAUAGUUUGCUUUAUGAGUAUUUUACGGUUUAUAACGAAUUGACAAAGGUCAAAUAUGUUACU<br>GAAGGAAUGCGAAAACCAGCAUUUCUUUCAGGUGAACAGAAGAAAGCCAUUGUUGAUUUACUCUUCAAAACAAAUCGAAAAGUAACCGUUAAGCAAUUAAAAGAAGA<br>UUAUUUCAAAAAAAUAGAAUGUUUUGAUAGUGUUGAAAUUUCAGGAGUUGAAGAUAGAUUUAAUGCUUCAUUAGGUACCUACCAUGAUUUGCUAAAAAUUAUUAAA<br>GAUAAAGAUUUUUUGGAUAAUGAAGAAAAUGAAGAUAUCUUAGAGGAUAUUGUUUUAACAUUGACCUUAUUUGAAGAUAGGGAGAUGAUUGAGGAAAGACUUAAAA<br>CAUAUGCUCACCUCUUUGAUGAUAAGGUGAUGAAACAGCUUAAACGUCGCCGUUAUACUGGUUGGGGACGUUUGUCUCGAAAAUUGAUUAAUGGUAUUAGGGAUAA<br>GCAAUCUGGCAAAACAAUAUUAGAUUUUUUGAAAUCAGAUGGUUUUGCCAAUCGCAAUUUUAUGCAGCUGAUCCAUGAUGAUAGUUUGACAUUUAAAGAAGACAUU<br>CAAAAAGCACAAGUGUCUGGACAAGGCGAUAGUUUACAUGAACAUAUUGCAAAUUUAGCUGGUAGCCCUGCUAUUAAAAAAGGUAUUUUACAGACUGUAAAAGUUG<br>UUGAUGAAUUGGUCAAAGUAAUGGGGCGGCAUAAGCCAGAAAAUAUCGUUAUUGAAAUGGCACGUGAAAAUCAGACAACUCAAAAGGGCCAGAAAAAUUCGCGAGA<br>GCGUAUGAAACGAAUCGAAGAAGGUAUCAAAGAAUUAGGAAGUCAGAUUCUUAAAGAGCAUCCUGUUGAAAAUACUCAAUUGCAAAAUGAAAAGCUCUAUCUCUAUU<br>AUCUCCAAAAUGGAAGAGACAUGUAUGUGGACCAAGAAUUAGAUAUUAAUCGUUUAAGUGAUUAUGAUGUCGAUCACAUUGUUCCACAAAGUUUCCUUAAAGACGA<br>UUCAAUAGACAAUAAGGUCUUAACGCGUUCUGAUAAAAAUCGUGGUAAAUCGGAUAACGUUCCAAGUGAAGAAGUAGUCAAAAAGAUGAAAAACUAUUGGAGACAAC<br>UUCUAAACGCCAAGUUAAUCACUCAACGUAAGUUUGAUAAUUUAACGAAAGCUGAACGUGGAGGUUUGAGUGAACUUGAUAAAGCUGGUUUUAUCAAACGCCAAUU<br>GGUUGAAACUCGCCAAAUCACUAAGCAUGUGGCACAAAUUUUGGAUAGUCGCAUGAAUACUAAAUACGAUGAAAAUGAUAAACUUAUUCGAGAGGUUAAAGUGAUU<br>ACCUUAAAAUCUAAAUUAGUUUCUGACUUCCGAAAAGAUUUCCAAUUCUAUAAAGUACGUGAGAUUAACAAUUACCAUCAUGCCCAUGAUGCGUAUCUAAAUGCCGU<br>CGUUGGAACUGCUUUGAUUAAGAAAUAUCCAAAACUUGAAUCGGAGUUUGUCUAUGGUGAUUAUAAAGUUUAUGAUGUUCGUAAAAUGAUUGCUAAGUCUGAGCAA<br>GAAAUAGGCAAAGCAACCGCAAAAUAUUUCUUUUACUCUAAUAUCAUGAACUUCUUCAAAACAGAAAUUACACUUGCAAAUGGAGAGAUUCGCAAACGCCCUCUAAU<br>CGAAACUAAUGGGGAAACUGGAGAAAUUGUCUGGGAUAAAGGGCGAGAUUUUGCCACAGUGCGCAAAGUAUUGUCCAUGCCCCAAGUCAAUAUUGUCAAGAAAACA<br>GAAGUACAGACAGGCGGAUUCUCCAAGGAGUCAAUUUUACCAAAAAGAAAUUCGGACAAGCUUAUUGCUCGUAAAAAAGACUGGGAUCCAAAAAAAUAUGGUGGUU<br>UUGAUAGUCCAACGGUAGCUUAUUCAGUCCUAGUGGUUGCUAAGGUGGAAAAAGGGAAAUCGAAGAAGUUAAAAUCCGUUAAAGAGUUACUAGGGAUCACAAUUAU<br>GGAAAGAAGUUCCUUUGAAAAAAAUCCGAUUGACUUUUUAGAAGCUAAAGGAUAUAAGGAAGUUAAAAAAGACUUAAUCAUUAAACUACCUAAAUAUAGUCUUUUU<br>GAGUUAGAAAACGGUCGUAAACGGAUGCUGGCUAGUGCCGGAGAAUUACAAAAAGGAAAUGAGCUGGCUCUGCCAAGCAAAUAUGUGAAUUUUUUAUAUUUAGCUA<br>GUCAUUAUGAAAAGUUGAAGGGUAGUCCAGAAGAUAACGAACAAAAACAAUUGUUUGUGGAGCAGCAUAAGCAUUAUUUAGAUGAGAUUAUUGAGCAAAUCAGUGA<br>AUUUUCUAAGCGUGUUAUUUUAGCAGAUGCCAAUUUAGAUAAAGUUCUUAGUGCAUAUAACAAACAUAGAGACAAACCAAUACGUGAACAAGCAGAAAAUAUUAUUC<br>AUUUAUUUACGUUGACGAAUCUUGGAGCUCCCGCUGCUUUUAAAUAUUUUGAUACAACAAUUGAUCGUAAACGAUAUACGUCUACAAAAGAAGUUUUAGAUGCCAC<br>UCUUAUCCAUCAAUCCAUCACUGGUCUUUAUGAAACACGCAUUGAUUUGAGUCAGCUAGGAGGUGACUGA |

#### Supplementary Table 4

| Name | Sequence |
|---|---|
| HA-WT | ATGAAAGCCATCATCGTGCTGCTGATGGTGGTGACATCTAATGCCGATAGAATCTGTACCGGCA<br>TCACCTCCTCTAACAGCCCTCACGTGGTCAAGACCGCCACACAGGGCGAGGTGAACGTGACCG<br>GCGTGATCCCCCTTACAACCACACCTACAAAAAGCCACTTCGCCAACCTGAAAGGAACCGAAA<br>CCAGAGGCAAACTGTGCCCCAAGTGCCTGAACTGCACCGACCTGGACGTGGCCCTGGGCAGA<br>CCTAAGTGTACCGGCAAGATCCCTAGCGCCAGAGTGTCTATCCTGCACGAGGTCAGACCTGTG<br>ACCTCCGGCTGCTTCCCCATCATGCACGACCGGACCAAGATCAGACAGCTGCCTAACCTGCTC<br>CGGGGATATGAACATGTGCGGCTGAGCACCCACAACGTGATTAACGCCGAGGACGCCCCTGG<br>CCGGCCTTACGAGATCGGCACCAGCGGCAGCTGTCCCAACATCACCAACGGCAATGGCTTCTT<br>CGCCACAATGGCCTGGGCCGTACCTAAGAACAAGACAGCTACAAACCCCCTGACCATCGAGGT<br>GCCTTATATCTGCACCGAAGGCGAAGACCAGATCACAGTGTGGGGCTTCCACAGCGACAGCGA<br>AACCCAGATGGCTAAACTGTACGGCGATAGCAAGCCCCAGAAATTCACCAGCTCCGCCAATGG<br>AGTGACAACACACTACGTGTCACAGATCGGCGGCTTTCCAAATCAAACAGAAGATGGAGGACT<br>GCCCCAGAGCGGCAGGATCGTGGTTGACTACATGGTGCAGAAGTCCGGCAAGACCGGCACCA<br>TCACCTACCAGAGGGGCATTCTGCTGCCACAAAAGGTGTGGTGCGCCAGCGGCAGAAGCAAG<br>GTGATCAAGGGATCTCTGCCTCTGATCGGAGAGGCCGATTGCCTGCATGAGAAGTACGGTGGC<br>CTGAACAAGAGCAAACCTTACTACACAGGAGAGCACGCCAAGGCTATTGGAAACTGCCCTATC<br>TGGGTGAAGACCCCTCTGAAGCTGGCTAACGGCACCAAGTACAGACCTCCAGCCAAGCTGCTG<br>AAGGAAAGAGGCTTCTTCGGCGCTATCGCCGGCTTCCTGGAAGGCGGATGGGAGGGGATGAT<br>CGCCGGATGGCACGGCTACACCTCTCACGGCGCCCACGGCGTGGCTGTGGCCGCAGATCTGA<br>AGTCTACACAGGAGGCCATCAACAAGATCACTAAGAATCTGAATAGCCTGTCTGAACTGGAAG<br>TGAAGAACCTGCAGCGGCTGTCCGGCGCCATGGACGAGCTGCACAATGAGATCCTGGAACTC<br>GACGAGAAAGTGGACGATCTGAGAGCCGACACCATCAGCAGCCAGATCGAGCTGGCCGTGCT<br>GCTGAGCAACGAGGGCATCATCAACAGCGAGGATGAGCACCTGCTGGCTCTGGAGAGAAAGC<br>TGAAAAAGATGCTGGGCCCCAGCGCCGTCGAGATCGGCAACGGATGTTTTGAGACAAAGCAC<br>AAGTGCAACCAGACCTGTCTGGACCGGATCGCCGCCGGCACATTCGACGCCGGCGAATTCAGC<br>CTGCCCACATTTGACAGCCTGAACATCACCGCCGCTTCTCTTAACGACGACGGCCTGGATAACC<br>ACACCATCCTGCTGTACTACAGCACCGCTGCTAGCAGCCTGGCCGTGACCCTGATGATCGCCAT<br>CTTCGTGGTGTACATGGTCTCTAGAGATAACGTGTCCTGCAGCATCTGCCTG |
| HA-M1 | ATGAAGGCCATCATCGTGCTGCTGATGGTGGTGACCAGCAACGCCGACCGGATCTGCACCGGG<br>ATCACCAGCAGCAACAGCCCCCACGTGGTGAAGACCGCCACCCAGGGGGAGGTGAACGTGAC<br>CGGGGTGATCCCCCTGACCACCACCCCCACCAAGAGCCACTTCGCCAACCTGAAGGGGACCG<br>AGACCCGGGGGAAGCTGTGCCCCAAGTGCCTGAACTGCACCGACCTGGACGTGGCCCTGGGG<br>CGGCCCAAGTGCACCGGGAAGATCCCCAGCGCCCGGGTGAGCATCCTGCACGAGGTGCGGCC<br>CGTGACCAGCGGGTGCTTCCCCATCATGCACGACCGGACCAAGATCCGGCAGCTGCCCAACCT<br>GCTGCGGGGGTACGAGCACGTGCGGCTGAGCACCCACAACGTGATCAACGCCGAGGACGCCC<br>CCGGGCGGCCCTACGAGATCGGGACCAGCGGGAGCTGCCCCAACATCACCAACGGGAACGG<br>GTTCTTCGCCACCATGGCCTGGGCCGTGCCCAAGAACAAGACCGCCACCAACCCCCTGACCAT<br>CGAGGTGCCCTACATCTGCACCGAGGGGGAGGACCAGATCACCGTGTGGGGGTTCCACAGCG<br>ACAGCGAGACCCAGATGGCCAAGCTGTACGGGGACAGCAAGCCCCAGAAGTTCACCAGCAGC<br>GCCAACGGGGTGACCACCCACTACGTGAGCCAGATCGGGGGGTTCCCCAACCAGACCGAGGA<br>CGGGGGGCTGCCCCAGAGCGGGCGGATCGTGGTGGACTACATGGTGCAGAAGAGCGGGAAG |

| | |
|---|---|
| | ACCGGGACCATCACCTACCAGCGGGGGATCCTGCTGCCCCAGAAGGTGTGGTGCGCCAGCGG<br>GCGGAGCAAGGTGATCAAGGGGAGCCTGCCCCTGATCGGGGAGGCCGACTGCCTGCACGAGA<br>AGTACGGGGGGCTGAACAAGAGCAAGCCCTACTACACCGGGGAGCACGCCAAGGCCATCGG<br>GAACTGCCCCATCTGGGTGAAGACCCCCCTGAAGCTGGCCAACGGGACCAAGTACCGGCCCC<br>CCGCCAAGCTGCTGAAGGAGCGGGGGTTCTTCGGGGCCATCGCCGGGTTCCTGGAGGGGGGG<br>TGGGAGGGGATGATCGCCGGGTGGCACGGGTACACCAGCCACGGGGCCCACGGGGTGGCCG<br>TGGCCGCCGACCTGAAGAGCACCCAGGAGGCCATCAACAAGATCACCAAGAACCTGAACAGC<br>CTGAGCGAGCTGGAGGTGAAGAACCTGCAGCGGCTGAGCGGGGCCATGGACGAGCTGCACAA<br>CGAGATCCTGGAGCTGGACGAGAAGGTGGACGACCTGCGGGCCGACACCATCAGCAGCCAGA<br>TCGAGCTGGCCGTGCTGCTGAGCAACGAGGGGATCATCAACAGCGAGGACGAGCACCTGCTG<br>GCCCTGGAGCGGAAGCTGAAGAAGATGCTGGGGCCCAGCGCCGTGGAGATCGGGAACGGGT<br>GCTTCGAGACCAAGCACAAGTGCAACCAGACCTGCCTGGACCGGATCGCCGCCGGGACCTTC<br>GACGCCGGGGAGTTCAGCCTGCCCACCTTCGACAGCCTGAACATCACCGCCGCCAGCCTGAAC<br>GACGACGGGCTGGACAACCACACCATCCTGCTGTACTACAGCACCGCCGCCAGCAGCCTGGC<br>CGTGACCCTGATGATCGCCATCTTCGTGGTGTACATGGTGAGCCGGGACAACGTGAGCTGCAG<br>CATCTGCCTG |
| HA-M2 | ATGAAGGCCATCATCGTGCTCCTCATGGTGGTGACGTCGAACGCCGACCGGATCTGCACGGGG<br>ATCACGTCGTCGAACTCGCCGCACGTGGTGAAGACGGCCACGCAGGGGGAGGTGAACGTGAC<br>GGGGGTGATCCCGCTCACGACGACGCCGACGAAGTCGCACTTCGCCAACCTCAAGGGGACGG<br>AGACGCGGGGGAAGCTCTGCCCGAAGTGCCTCAACTGCACGGACCTCGACGTGGCCCTCGGG<br>CGGCCGAAGTGCACGGGGAAGATCCCGTCGGCCCGGGTGTCGATCCTCCACGAGGTGCGGCC<br>GGTGACGTCGGGGTGCTTCCCGATCATGCACGACCGGACGAAGATCCGGCAGCTCCCGAACCT<br>CCTCCGGGGGTACGAGCACGTGCGGCTCTCGACGCACAACGTGATCAACGCCGAGGACGCCC<br>CGGGGCGGCCGTACGAGATCGGGACGTCGGGGTCGTGCCCGAACATCACGAACGGGAACGG<br>GTTCTTCGCCACGATGGCCTGGGCCGTGCCGAAGAACAAGACGGCCACGAACCCGCTCACGAT<br>CGAGGTGCCGTACATCTGCACGGAGGGGGAGGACCAGATCACGGTGTGGGGGTTCCACTCGG<br>ACTCGGAGACGCAGATGGCCAAGCTCTACGGGGACTCGAAGCCGCAGAAGTTCACGTCGTCG<br>GCCAACGGGGTGACGACGCACTACGTGTCGCAGATCGGGGGGTTCCCGAACCAGACGGAGGA<br>CGGGGGGCTCCCGCAGTCGGGGCGGATCGTGGTGGACTACATGGTGCAGAAGTCGGGGAAGA<br>CGGGGACGATCACGTACCAGCGGGGGATCCTCCTCCCGCAGAAGGTGTGGTGCGCCTCGGGG<br>CGGTCGAAGGTGATCAAGGGGTCGCTCCCGCTCATCGGGGAGGCCGACTGCCTCCACGAGAA<br>GTACGGGGGGCTCAACAAGTCGAAGCCGTACTACACGGGGGAGCACGCCAAGGCCATCGGGA<br>ACTGCCCGATCTGGGTGAAGACGCCGCTCAAGCTCGCCAACGGGACGAAGTACCGGCCGCCG<br>GCCAAGCTCCTCAAGGAGCGGGGGTTCTTCGGGGCCATCGCCGGGTTCCTCGAGGGGGGGTG<br>GGAGGGGATGATCGCCGGGTGGCACGGGTACACGTCGCACGGGGCCCACGGGGTGGCCGTG<br>GCCGCCGACCTCAAGTCGACGCAGGAGGCCATCAACAAGATCACGAAGAACCTCAACTCGCT<br>CTCGGAGCTCGAGGTGAAGAACCTCCAGCGGCTCTCGGGGGCCATGGACGAGCTCCACAACG<br>AGATCCTCGAGCTCGACGAGAAGGTGGACGACCTCCGGGCCGACACGATCTCGTCGCAGATC<br>GAGCTCGCCGTGCTCCTCTCGAACGAGGGGATCATCAACTCGGAGGACGAGCACCTCCTCGCC<br>CTCGAGCGGAAGCTCAAGAAGATGCTCGGGCCGTCGGCCGTGGAGATCGGGAACGGGTGCTT<br>CGAGACGAAGCACAAGTGCAACCAGACGTGCCTCGACCGGATCGCCGCCGGGACGTTCGACG<br>CCGGGGAGTTCTCGCTCCCGACGTTCGACTCGCTCAACATCACGGCCGCCTCGCTCAACGACG<br>ACGGGCTCGACAACCACACGATCCTCCTCTACTACTCGACGGCCGCCTCGTCGCTCGCCGTGA |

| | |
|---|---|
| | CGCTCATGATCGCCATCTTCGTGGTGTACATGGTGTCGCGGGACAACGTGTCGTGCTCGATCTG<br>CCTC |
| HA-M3 | ATGAAGGCCATCATCGTTCTGCTGATGGTGGTGACCAGCAATGCGGACCGGATATGTACTGGT<br>ATTACCTCTAGTAACAGTCCCCATGTGGTGAAGACTGCGACTCAGGGGGAGGTGAACGTCACG<br>GGGGTGATCCCCCTGACGACCACCCCCACCAAGTCGCACTTCGCCAACCTCAAGGGGACTGAG<br>ACTAGAGGTAAGCTTTGTCCGAAGTGCCTGAACTGTACCGATCTCGACGTGGCGCTGGGCCGG<br>CCCAAGTGCACTGGGAAGATTCCCAGTGCACGGGTGAGCATCTTGCATGAGGTGCGCCCAGTG<br>ACCAGCGGGTGCTTTCCCATTATGCATGATCGGACAAAGATTCGGCAGCTGCCGAATCTCCTCC<br>GCGGGTATGAGCATGTGCGTCTGTCGACGCACAATGTCATAAACGCGGAGGACGCTCCCGGAC<br>GTCCCTATGAGATAGGGACGTCCGGGAGCTGTCCGAACATCACTAATGGGAACGGGTTCTTCG<br>CCACCATGGCTTGGGCCGTGCCCAAGAACAAGACGGCGACGAACCCCCTCACTATTGAGGTGC<br>CCTACATCTGTACTGAGGGGGAGGACCAGATCACCGTGTGGGGCTTCCATTCAGACTCCGAGA<br>CTCAGATGGCAAAGCTCTATGGCGACAGTAAGCCGCAGAAGTTCACCTCCTCTGCCAACGGTG<br>TGACCACCCACTATGTGAGCCAGATTGGGGGGTTCCCCAACCAGACCGAGGATGGGGGACTC<br>CCCCAGTCTGGCCGCATAGTGGTGGATTACATGGTGCAGAAGAGCGGCAAGACGGGCACCAT<br>CACCTATCAGCGGGGGATCCTTCTCCCGCAGAAGGTGTGGTGCGCGTCTGGCCGCTCTAAGGT<br>GATTAAGGGGAGTCTTCCCCTGATCGGGGAGGCCGATTGCCTTCACGAGAAGTACGGCGGGCT<br>CAACAAGAGCAAGCCGTACTACACTGGTGAGCACGCCAAGGCAATCGGCAACTGCCCGATCT<br>GGGTGAAGACTCCCCTTAAGCTGGCGAACGGTACCAAGTACCGGCCGCCAGCTAAGCTGCTCA<br>AGGAGCGGGGGTTCTTCGGGGCCATCGCTGGGTTTCTTGAGGGCGGCTGGGAGGGCATGATC<br>GCTGGGTGGCATGGCTACACTAGCCATGGCGCCCACGGTGTTGCCGTCGCAGCCGACCTCAAG<br>AGCACCCAGGAGGCCATCAACAAGATCACGAAGAACCTCAACTCCTTGAGCGAGCTGGAGGT<br>GAAGAACCTTCAGCGGCTGTCTGGCGCCATGGACGAGTTGCACAATGAGATCTTGGAGCTGGA<br>TGAGAAGGTCGACGATCTGAGAGCTGATACCATCAGCTCTCAGATCGAGCTGGCCGTCCTCCT<br>CAGTAATGAGGGCATCATCAATAGTGAGGACGAGCACCTGCTGGCACTGGAGCGCAAGCTCA<br>AGAAGATGCTCGGGCCCAGCGCCGTCGAGATCGGTAACGGGTGCTTCGAGACAAAGCACAAG<br>TGCAACCAGACATGTCTGGACCGCATTGCTGCAGGGACGTTCGACGCGGGAGAGTTCTCTCTC<br>CCGACGTTCGACTCCCTGAACATCACCGCCGCCAGCCTGAACGATGATGGCCTTGATAACCAC<br>ACAATATTGCTGTATTACAGCACCGCCGCCAGTTCTCTGGCGGTGACGCTGATGATAGCAATAT<br>TTGTGGTCTACATGGTGAGTAGAGACAACGTGAGTTGCTCTATTTGCCTG |
| HA-M4 | ATGAAGGCCATCATCGTTCTGCTCATGGTGGTGACCAGCAATGCGGACCGGATATGTACTGGTA<br>TTACCTCGAGTAACAGTCCCCATGTGGTGAAGACTGCGACTCAGGGGGAGGTCAACGTCACGG<br>GGGTGATCCCCCTGACGACGACCCCCACCAAGTCGCACTTCGCCAACCTCAAGGGGACTGAGA<br>CTCGAGGTAAGCTTTGTCCGAAGTGCCTGAACTGTACCGATCTCGACGTGGCGCTGGGCCGAC<br>CCAAGTGCACTGGGAAGATTCCCAGTGCACGGGTCTCCATCCTCCACGAGGTTCGGCCGGTCA<br>CGAGTGGGTGCTTTCCGATCATGCATGACCGCACGAAGATTCGGCAGCTGCCGAATCTTCTGC<br>GCGGTTATGAGCATGTTCGGCTGAGCACCCACAACGTGATCAACGCCGAGGACGCTCCCGGAC<br>GTCCCTATGAGATAGGGACGTCCGGGAGCTGTCCCAACATCACCAATGGCAACGGGTTCTTCG<br>CCACCATGGCTTGGGCCGTGCCCAAGAACAAGACGGCGACGAACCCGTTGACCATTGAGGTTC<br>CGTACATCTGTACGGAAGGGGAGGACCAGATTACTGTCTGGGGCTTCCATAGTGACTCCGAGA<br>CGCAGATGGCGAAGCTTTATGGGGACAGCAAGCCCCAGAAGTTTACGTCATCTGCGAACGGAG<br>TCACTACCCACTATGTGAGCCAGATTGGGGGGTTCCCCAACCAGACCGAGGATGGGGGACTCC<br>CCCAGTCTGGCCGCATAGTGGTGGACTACATGGTCCAGAAGAGCGGCAAGACGGGCACCATC |

| | |
|---|---|
| | ACCTATCAGCGGGGGATCCTTCTCCCGCAGAAGGTGTGGTGCGCGTCTGGCCGCTCTAAGGTG<br>ATTAAGGGGAGTCTTCCCCTGATCGGGGAGGCCGATTGCCTTCACGAGAAGTACGGCGGGCTC<br>AACAAGAGCAAGCCGTACTACACTGGTGAGCACGCCAAGGCAATCGGCAACTGCCCGATCTG<br>GGTGAAGACTCCCCTTAAGCTGGCGAACGGTACCAAGTACCGTCCGCCAGCTAAGTTGCTCAA<br>GGAGCGGGGGTTCTTCGGGGCCATCGCTGGGTTTCTTGAGGGCGGCTGGGAGGGCATGATCG<br>CGGGGTGGCATGGCTACACTAGCCATGGCGCCCACGGGGTCGCCGTCGCAGCCGACCTCAAG<br>AGCACCCAGGAGGCCATCAACAAGATCACGAAGAACCTCAACTCCTTGAGCGAGCTGGAGGT<br>GAAGAACCTCCAGCGCCTCAGTGGAGCGATGGACGAGTTGCATAATGAGATCCTCGAGCTCGA<br>CGAGAAGGTCGACGATCTGCGAGCTGATACCATCAGCTCGCAGATCGAGTTGGCCGTCTTGTT<br>GAGCAACGAGGGCATTATCAACTCGGAGGACGAGCATCTTCTTGCGCTTGAGCGCAAGCTCAA<br>GAAGATGCTCGGGCCCAGCGCCGTCGAGATCGGTAACGGGTGCTTCGAGACAAAGCACAAGT<br>GCAACCAGACATGTCTGGACCGCATTGCTGCAGGGACGTTCGACGCGGGAGAGTTCTCTCTCC<br>CGACGTTCGACTCCCTGAACATCACCGCCGCGAGCCTGAACGATGATGGCCTTGATAACCACA<br>CGATATTGCTGTATTACAGCACCGCCGCCAGTTCTCTGGCGGTGACGCTGATGATAGCAATATT<br>TGTGGTCTACATGGTAAGTCGAGACAACGTGAGTTGCTCGATTTGCCTG |

#### Supplementary Table 5

The similarity between HA-WT and M1,M2,M3,M4:

| **Sequence** | **HA-M1** | **HA-M2** | **HA-M3** | **HA-M4** |
|---|---|---|---|---|
| **Similarity** | 0.6701030927835051 | 0.4742268041237113 | 0.47079037800687284 | 0.45017182130584193 |

#### Supplementary Table 6

| **Name** | **Length** | **MEF** | **CAI** | **Similarity to WT Bmp2** |
|---|---|---|---|---|
| WT Bmp2 | 1251 | -420.900 | 0.770 | 1.000000 |
| GEMORNA | 1251 | -447.200 | 0.789 | 0.470024 |
| RNAjog | 1251 | -522.300 | 0.990 | 0.494005 |
| RNAjog-m6A-1 | 1251 | -529.900 | 0.981 | 0.486811 |
| RNAjog-m6A-2 | 1251 | -529.200 | 0.976 | 0.484412 |

| **Name** | **Sequence** |
|---|---|
| WT Bmp2 | ATGGTGGCCGGGACCCGCTGTCTTCTAGTGTTGCTGCTTCCCCAGGTCCTCC<br>TGGGCGGCGCGGCCGGCCTCATTCCAGAGCTGGGCCGCAAGAAGTTCGCC<br>GCGGCATCCAGCCGACCCTTGTCCCGGCCTTCGGAAGACGTCCTCAGCGAA<br>TTTGAGTTGAGGCTGCTCAGCATGTTTGGCCTGAAGCAGAGACCCACCCCCA<br>GCAAGGACGTCGTGGTGCCCCCCTATATGCTAGATCTGTACCGCAGGCACT<br>CAGGCCAGCCAGGAGCGCCCGCCCCAGACCACCGGCTGGAGAGGGCAGC<br>CAGCCGCGCCAACACCGTGCGCAGCTTCCATCACGAAGAAGCCGTGGAGG |

| | |
|---|---|
| | AACTTCCAGAGATGAGTGGGAAAACGGCCCGGCGCTTCTTCTTCAATTTAAG<br>TTCTGTCCCCAGTGACGAGTTTCTCACATCTGCAGAACTCCAGATCTTCCGG<br>GAACAGATACAGGAAGCTTTGGGAAACAGTAGTTTCCAGCACCGAATTAAT<br>ATTTATGAAATTATAAAGCCTGCAGCAGCCAACTTGAAATTTCCTGTGACCA<br>GACTATTGGACACCAGGTTAGTGAATCAGAACACAAGTCAGTGGGAGAGCT<br>TCGACGTCACCCCAGCTGTGATGCGGTGGACCACACAGGGACACACCAACC<br>ATGGGTTTGTGGTGGAAGTGGCCCATTTAGAGGAGAACCCAGGTGTCTCCA<br>AGAGACATGTGAGGATTAGCAGGTCTTTGCACCAAGATGAACACAGCTGGT<br>CACAGATAAGGCCATTGCTAGTGACTTTTGGACATGATGGAAAAGGACATCC<br>GCTCCACAAACGAGAAAAGCGTCAAGCCAAACACAAACAGCGGAAGCGCC<br>TCAAGTCCAGCTGCAAGAGACACCCTTTGTATGTGGACTTCAGTGATGTGGG<br>GTGGAATGACTGGATCGTGGCACCTCCGGGCTATCATGCCTTTTACTGCCAT<br>GGGGAGTGTCCTTTTCCCCTTGCTGACCACCTGAACTCCACTAACCATGCCA<br>TAGTGCAGACTCTGGTGAACTCTGTGAATTCCAAAATCCCTAAGGCATGCTG<br>TGTCCCCACAGAGCTCAGCGCAATCTCCATGTTGTACCTAGATGAAAATGAA<br>AAGGTTGTGCTAAAAAATTATCAGGACATGGTTGTGGAGGGCTGCGGGTGT<br>CGTGGTGGTGGCTCTGGTGGCGGCGGTTCTGGTGGCGGCGGCTACCCCTAC<br>GATGTGCCCGACTACGCCTAG |
| GEMORNA | ATGGTGGCGGGCACCCGCTGCCTGCTGGTGCTGCTGCTGCCCCAGGTGCTG<br>CTGGGCGGGGCGGCGGGGCTGATCCCGGAGCTGGGCCGGAAGAAGTTCGC<br>GGCGGCCTCGAGCCGCCCCCTGTCTCGGCCGTCGGAGGACGTGCTGAGCG<br>AGTTCGAGCTGCGCCTGCTGTCCATGTTCGGCCTGAAGCAGCGGCCCACCC<br>CCAGCAAGGACGTGGTGGTCCCCCCGTACATGCTGGACCTGTACCGCAGGC<br>ACAGCGGCCAGCCCGGGGCGCCGGCTCCGGACCACCGCCTGGAGCGCGCC<br>GCCAGCAGGGCCAACACGGTCCGCAGCTTCCACCACGAGGAGGCGGTGGA<br>GGAGCTGCCCGAGATGTCCGGCAAGACGGCCCGCCGCTTCTTCTTCAACCT<br>GTCCTCCGTGCCCAGCGACGAGTTCCTGACCTCGGCCGAGCTGCAGATCTTC<br>CGCGAGCAGATCCAGGAGGCGCTGGGCAACAGCAGCTTCCAGCACCGCAT<br>CAACATTTATGAAATTATTAAACCTGCTGCTGCTAATCTTAAGTTTCCTGTCAC<br>TAGACTTCTTGATACAAGACTTGTCAATCAGAACACATCTCAGTGGGAATCTT<br>TTGATGTAACTCCTGCAGTAATGAGATGGACCACCCAGGGACATACTAACCA<br>TGGGTTTGTTGTTGAAGTTGCTCATCTTGAAGAAAATCCTGGTGTCAGCAAAA<br>GACATGTTAGAATTTCTAGAAGTCTTCATCAAGATGAACATAGCTGGAGTCA<br>GATTCGACCTCTTCTTGTGACTTTTGGTCATGATGGAAAAGGTCATCCTCTTC<br>ATAAAAGAGAAAAAAGACAAGCTAAACATAAACAGAGAAAAAGGCTTAAAT<br>CTTCATGTAAAAGACATCCACTGTATGTTGATTTTTCTGATGTTGGATGGAAT<br>GACTGGATTGTCGCACCTCCTGGATATCATGCTTTTTACTGTCATGGAGAATG<br>TCCATTTCCTTTAGCAGATCATCTTAATAGTACAAATCATGCTATTGTGCAGA<br>CTTTGGTGAATTCCGTGAACAGCAAGATTCCCAAAGCCTGCTGCGTCCCAAC<br>AGAGCTGTCCGCCATCTCCATGCTCTATCTGGATGAGAATGAGAAGGTGGTG<br>CTTAAGAATTACCAGGACATGGTGGTGGAGGGCTGTGGATGTAGGGGTGGT<br>GGCTCTGGTGGCGGCGGTTCTGGTGGCGGCGGCTACCCCTACGATGTGCCC<br>GACTACGCCTAG |
| RNAjog | ATGGTGGCCGGCACCCGGTGCCTGCTGGTGCTGCTGCTGCCTCAGGTGCTG |

| | |
|---|---|
| | CTGGGCGGCGCCGCCGGCCTGATCCCTGAGCTGGGCAGGAAGAAGTTCGC<br>CGCCGCCTCCTCCCGGCCTCTGAGCAGACCTTCTGAGGACGTGCTGTCCGA<br>GTTCGAGCTGAGACTGCTGAGCATGTTCGGCCTGAAGCAGAGACCTACCCC<br>TAGCAAGGACGTGGTGGTGCCCCCCTACATGCTGGACCTGTACCGGCGGCA<br>CAGCGGCCAGCCTGGCGCCCCTGCCCCTGACCACCGGCTGGAGCGGGCCG<br>CCTCTCGGGCCAACACCGTGAGAAGCTTCCACCACGAGGAGGCCGTGGAG<br>GAGCTGCCTGAGATGAGCGGCAAGACCGCCCGGCGGTTCTTCTTCAACCTG<br>AGCAGCGTGCCCAGCGACGAGTTCCTGACCAGCGCCGAGCTGCAGATCTTC<br>AGAGAGCAGATCCAGGAGGCCCTGGGCAACAGCAGCTTCCAGCACCGGAT<br>CAACATCTACGAGATCATCAAGCCTGCCGCCGCCAACCTGAAGTTCCCTGTG<br>ACCCGGCTGCTGGACACCCGGCTGGTGAACCAGAACACCAGCCAGTGGGA<br>GTCCTTCGACGTGACCCCCGCCGTGATGCGGTGGACCACCCAGGGCCACAC<br>CAACCACGGCTTCGTGGTGGAGGTGGCCCACCTGGAGGAGAACCCTGGCGT<br>GAGCAAGCGGCACGTGCGGATCAGCCGGAGCCTGCACCAGGACGAGCACA<br>GCTGGAGCCAGATCCGGCCTCTGCTGGTGACCTTCGGCCACGACGGCAAGG<br>GCCACCCTCTGCACAAGAGGGAGAAGAGGCAGGCCAAGCACAAGCAGCGG<br>AAGAGACTGAAGAGCAGCTGCAAGCGGCACCCTCTGTACGTGGACTTCAGC<br>GACGTGGGCTGGAACGACTGGATCGTGGCCCCTCCTGGCTACCACGCCTTC<br>TACTGCCACGGCGAGTGCCCTTTCCCTCTGGCCGACCACCTGAACAGCACC<br>AACCACGCCATCGTGCAGACCCTGGTGAACTCCGTGAACAGCAAGATCCCT<br>AAGGCCTGCTGCGTGCCAACCGAGCTGAGCGCCATCAGCATGCTGTACCTG<br>GACGAGAACGAGAAGGTGGTGCTGAAGAACTACCAGGACATGGTGGTGGA<br>GGGCTGCGGCTGCCGGGGTGGTGGCTCTGGTGGCGGCGGTTCTGGTGGCG<br>GCGGCTACCCCTACGATGTGCCCGACTACGCCTAG |
| RNAjog-<br>m6A-1 | ATGGTGGCCGGCACCCGGTGCCTGCTGGTGCTGCTGCTGCCTCAGGTGCTG<br>CTGGGCGGCGCCGCCGGCCTGATCCCTGAGCTGGGCCGGAAGAAGTTCGC<br>CGCCGCCAGCAGCCGGCCTCTGAGCCGGCCTAGCGAGGACGTGCTGAGCG<br>AGTTCGAGCTGCGGCTGCTGAGCATGTTCGGCCTGAAGCAGCGGCCTACCC<br>CTAGCAAGGACGTGGTGGTGCCTCCTTACATGCTGGATCTGTACCGGCGGC<br>ACAGCGGCCAGCCTGGCGCCCCTGCCCCTGACCACCGGCTGGAGCGGGCC<br>GCCAGCCGGGCCAACACCGTGCGGAGCTTCCACCACGAGGAGGCCGTGGA<br>GGAGCTGCCTGAGATGAGCGGCAAGACCGCCCGGCGGTTCTTCTTCAACCT<br>GAGCAGCGTGCCTAGCGACGAGTTCCTGACCAGCGCCGAGCTGCAGATCTT<br>CCGGGAGCAGATCCAGGAGGCCCTGGGCAACAGCAGCTTCCAGCACCGGA<br>TCAACATCTACGAGATCATCAAGCCTGCCGCCGCCAACCTGAAGTTCCCTGT<br>GACCCGGCTGCTGGATACCCGGCTGGTGAACCAGAATACCAGCCAGTGGG<br>AGAGCTTCGACGTGACCCCTGCCGTGATGCGGTGGACGACCCAGGGCCACA<br>CCAACCACGGCTTCGTGGTGGAGGTGGCCCACCTGGAGGAGAACCCTGGC<br>GTGAGCAAGCGGCACGTGCGGATCAGCCGGAGCCTGCACCAGGACGAGCA<br>CAGCTGGAGCCAGATCCGGCCTCTGCTGGTGACCTTCGGCCACGACGGCAA<br>GGGCCACCCTCTGCACAAGCGGGAGAAGCGGCAGGCCAAGCACAAGCAGC<br>GGAAGCGGCTGAAGAGCAGCTGCAAGCGGCACCCTCTGTACGTGGATTTCA<br>GCGACGTGGGCTGGAACGACTGGATCGTGGCCCCTCCTGGCTACCACGCCT<br>TCTACTGCCACGGCGAGTGCCCTTTCCCTCTGGCCGACCACCTGAATAGCAC |

| | |
|---|---|
| | CAACCACGCCATCGTGCAGACCCTGGTGAATAGCGTGAATAGCAAGATCCC<br>TAAGGCCTGCTGCGTGCCTACCGAGCTGAGCGCCATCAGCATGCTGTACCT<br>GGACGAGAACGAGAAGGTGGTGCTGAAGAATTACCAGGATATGGTGGTGG<br>AGGGCTGCGGCTGCCGGGGTGGTGGCTCTGGTGGCGGCGGTTCTGGTGGC<br>GGCGGCTACCCCTACGATGTGCCCGACTACGCCTAG |
| RNAjog-m6A-2 | ATGGTGGCCGGCACCCGGTGCCTGCTGGTGCTGCTGCTGCCTCAGGTGCTG<br>CTGGGCGGCGCCGCCGGCCTGATCCCTGAGCTGGGCCGGAAGAAGTTCGC<br>CGCCGCCAGCAGCCGGCCTCTGAGCCGGCCTAGCGAGGACGTGCTGAGCG<br>AGTTCGAGCTGCGGCTGCTGAGCATGTTCGGCCTGAAGCAGCGGCCTACCC<br>CTAGCAAGGACGTGGTGGTGCCTCCTTACATGCTGGATCTGTACCGGCGGC<br>ACAGCGGCCAGCCTGGCGCCCCTGCCCCTGAtCACCGGCTGGAGCGGGCCG<br>CCAGCCGGGCCAACACCGTGCGGAGCTTCCACCACGAGGAGGCCGTGGAG<br>GAGCTGCCTGAGATGAGCGGCAAaACgGCCCGGCGGTTCTTCTTCAACCTG<br>AGCAGCGTGCCTAGCGACGAGTTCCTcACCAGCGCCGAGCTGCAGATCTTC<br>CGGGAGCAGATCCAGGAGGCCCTGGGCAACAGCAGCTTCCAGCACCGGAT<br>CAACATCTACGAGATCATCAAGCCTGCCGCCGCCAACCTGAAGTTCCCTGTG<br>ACCCGGCTGCTGGATACCCGGCTGGTGAAtCAGAATACCAGCCAGTGGGAG<br>AGCTTCGACGTcACCCCTGCCGTGATGCGGTGGACGACCCAGGGCCACACC<br>AACCACGGCTTCGTGGTGGAGGTGGCCCACCTGGAGGAGAACCCTGGCGT<br>GAGCAAGCGGCACGTGCGGATCAGCCGGAGCCTGCACCAGGACGAGCACA<br>GCTGGAGCCAGATCCGGCCTCTGCTGGTGACCTTCGGCCACGACGGCAAGG<br>GCCACCCTCTGCACAAGCGGGAGAAGCGGCAGGCCAAGCACAAGCAGCGG<br>AAGCGGCTGAAGAGCAGCTGCAAGCGGCACCCTCTGTACGTGGATTTCAGC<br>GACGTGGGCTGGAACGACTGGATCGTGGCCCCTCCTGGCTACCACGCCTTC<br>TACTGCCACGGCGAGTGCCCTTTCCCTCTGGCCGACCACCTGAATAGCACCA<br>ACCACGCCATCGTGCAGACCCTGGTGAATAGCGTGAATAGCAAGATCCCTA<br>AGGCCTGCTGCGTGCCTACCGAGCTGAGCGCCATCAGCATGCTGTACCTGG<br>ACGAGAACGAGAAGGTGGTGCTGAAGAATTACCAGGATATGGTGGTGGAG<br>GGCTGCGGCTGCCGGGGTGGTGGCTCTGGTGGCGGCGGTTCTGGTGGCGG<br>CGGCTACCCCTACGATGTGCCCGACTACGCCTAG |

### Supplementary Table 7

| Antibody Name | Catalogue No. | Company |
|---|---|---|
| Anti-HA tag antibody | ab182009 | abcam |
| Vinculin Polyclonal antibody | 26520-1-AP | proteintech |
| HRP-conjugated Goat Anti-Rabbit IgG(H+L) | SA00001-2 | proteintech |

### Supplementary Table 8

| Primer name | Sequence |
|---|---|
| KanR-F | GGGACTGGCTGCTATTGG |
| KanR-R | CGATGTTTCGCTTGGTGGT |
| Bmp2-F | GGTGGTGGCTCTGGTGGC |
| Bmp2-R | GGCTGATCAGCGGGTTTAAACG |
